# Dynamic changes in tRNA modifications and abundance during T-cell activation

**DOI:** 10.1101/2020.03.14.991901

**Authors:** Roni Rak, Michal Polonsky, Inbal Eizenberg-Magar, Yufeng Mo, Yuriko Sakaguchi, Orel Mizrahi, Aharon Nachshon, Shlomit Reich-Zeliger, Noam Stern-Ginossar, Orna Dahan, Tsutomu Suzuki, Nir Friedman, Yitzhak Pilpel

## Abstract

The tRNA pool determines the efficiency, throughput, and accuracy of translation. Previous studies have identified dynamic changes in the tRNA supply and mRNA demand during cancerous proliferation. Yet, dynamic changes may occur also during physiologically normal proliferation, and these are less characterized. We examined the tRNA and mRNA pools of T-cells during their vigorous proliferation and differentiation upon triggering their antigen receptor. We observe a global signature of switch in demand for codons at the early proliferation phase of the response, accompanied by corresponding changes in tRNA expression levels. In the later phase, upon differentiation, the response of the tRNA pool is relaxed back to basal level, potentially restraining excessive proliferation. Sequencing of tRNAs allowed us to also evaluate their diverse base-modifications. We found that two types of tRNA modifications, wybutosine and ms^2^t6A, are reduced dramatically during T-cell activation. These modifications occur in the anti-codon loops of two tRNAs that decode “slippery codons”, that are prone to ribosomal frameshifting. Attenuation of these frameshift-protective modifications is expected to increase the potential for proteome-wide frameshifting during T-cell proliferation. Indeed, human cell lines deleted of a wybutosine writer showed increased ribosomal frameshifting, as detected with a HIV gag-pol frameshifting site reporter. These results may explain HIV’s specific tropism towards proliferating T-Cells since it requires ribosomal frameshift exactly on the corresponding codon for infection. The changes in tRNA expression and modifications uncover a new layer of translation regulation during T-cell proliferation and exposes a potential trade-off between cellular growth and translation fidelity.

**Significance statement:** The tRNA pool decodes genetic information during translation. As such, it is subject to intricate physiological regulation in all species, across different physiological conditions. Here we show for the first time a program that governs the tRNA pool and its interaction with the transcriptome upon a physiological cellular proliferation- T-cells activation. We found that upon antigenic activation of T-cells, their tRNA and mRNA pools undergo coordinated and complementary changes, which are relaxed when cells reduce back their proliferation rate and differentiate into memory cells. We found a reduction in two particular tRNA modifications that have a role in governing translation fidelity and frameshift prevention. This exposes a vulnerability in activated T-cells that may be utilized by HIV for its replication.

**Classification:** BIOLOGICAL SCIENCES; cell biology

## Introduction

Few cells in the adult mammalian body can proliferate under normal conditions. One example of a fundamental programmed proliferation processes is evoked in clonal expansion of selected cells of the adaptive immune system, following encounter with foreign antigens. Upon recognition of a cognate antigen, naive T-cells (and B-cells) undergo massive proliferation and differentiation, hence changing their status from arrested naive cells to highly proliferating effector cells^1^. The integrity of the immune response is dependent on the precise regulation of proliferation rates, number of cell-cycles and cell death. In addition, a portion of the population is differentiated into memory cells, which remain in the body for extended periods of time. A major regulatory challenge and point of vulnerability is to allow such massive cellular proliferation while circumventing the risk of cancerous growth.

A crucial part of cell proliferation is the ability to massively translate new proteins. Translation control was shown to regulate gene expression in the immune system^2–4^. However, the regulation of translation elongation, and especially tRNA availability was not explored. We have previously shown that genes that are upregulated in proliferating cancerous cells or upon induced pluripotency have a distinct translation program from that of arrested cells ^5,6^. In particular, mRNAs corresponding to cell-autonomous functions, related to proliferating cells, are enriched with a specific set of codons, while mRNA of multicellular functions, related to non-dividing cells, are enriched with a different set of preferred codons. The tRNA pool in these cells is dynamically regulated and its dynamics corresponds to the changes in codon usage, as also reflected in differential tRNA essentiality in proliferation and cell arrest^7,8^. Following these observations, it is essential to explore how does the tRNA pool change in a natural proliferation process, and what are the contributions of the tRNA pool regulation to the integrity dynamics and of the immune response.

Beyond changes in expression level of tRNAs, these RNAs are among the most highly post-transcriptionally modified molecules in the mammalian transcriptome^9,10^ Essentially all tRNA molecules undergo diverse chemical modifications on many of their bases, and the various nucleotide modifications can affect tRNA folding and stability, amino acid loading, and codon-anticodon base pairing of tRNAs to their corresponding codons^11–15^. Modifications around the anticodon loop of the tRNA are known to affect translation fidelity, speed, and tendency to frameshift^15–18^. Modification levels are regulated and can change in a dynamic manner across physiological conditions^19–21^, in cancer^22,23^, and during development^24^. Yet the dynamics, regulation and biological function, both at the molecular and physiological level, of most such modifications are not well-characterized.

Here we set to investigate the control of the tRNA pool in a physiologically normal and programmed proliferation process, by following T-cells upon their antigen receptor activation. We examined the codon usage changes during T-cell activation and the corresponding changes in tRNA abundance. We also examined in parallel the post-transcriptional chemical base modifications of the tRNAs. We found that upon activation, T-cells reprogram their tRNA pool to serve the altered codon usage demand of the proliferation related genes. Interestingly though, at the later stage of the response, when T-cells differentiate, the tRNA response is relaxed back towards base level, where it is no longer adapted to the proliferation related codon usage. Further, we found that at the pick of their proliferation, T-cells exhibit a sharp decline of two tRNA base modifications. Both of these modifications are known to protect the ribosome from -1 frameshifting at “slippery” frameshift-prone codons. Interestingly, the reproduction of human immunodeficiency virus-1 (HIV-1), which preferentially targets proliferating T-cells, depends exactly on such -1 frameshifting that occurs on one of these slippery codons^25^. We indeed found that knockout of a key enzyme in one of these two tRNA modifications pathways, in human cells, induced a frameshift at the HIV frameshifting slippery codon sequence. While the biological reason for reduction of this modification in T-cells is still unknown, we hypothesize that it may represent a tendency to trade-off translation accuracy for speed, a potential requirement for fast proliferation.

## Results

### A global codon usage shift, towards AT-ending codon, followed by a relaxation during T-cell activation and differentiation

An important aspect of translation regulation is the dynamic change of supply (tRNA and anti-codons) and demand (mRNA and codons)^7^. To study translation regulatory dynamics during a central process of mammalian cellular proliferation and differentiation, we followed T-cells as they are triggered to switch from a naïve to an activated state. Naïve CD4+ T-cells were isolated from mice spleen and activated using anti-CD3/anti-CD28 activation beads (see Material and Methods). Samples were taken at the naïve stage (“time point zero”), and after 20, 48, and 72 hours. At the later time points (i.e. 48 and 72 hours) cells were sorted according to expression of the CD62L and CD44 markers. High CD62L and CD44 cells (CD62L^+^ CD44^+^) correspond with differentiation toward precursor central memory^26^, while CD44^+^, a marker of T-cells activation, was high in all activated cells (Fig.1A). The experiment was repeated twice. We sequenced both mRNAs and tRNAs from each sample. Unsorted samples from one biological repeat were analyzed also by ribosomal protection sequencing and LC/MS shotgun analyses (Fig 1A, see Material and Methods). By examining expression pattern of cell cycle genes, we found that 20h after activation cells are at the peak of S-phase gene expression, while the peak of M-phase gene expression is at 48h after activation (Fig. 1B). Those results are in line with the known dynamic of T-cells activation in-vitro^26^.

**Figure 1-.**
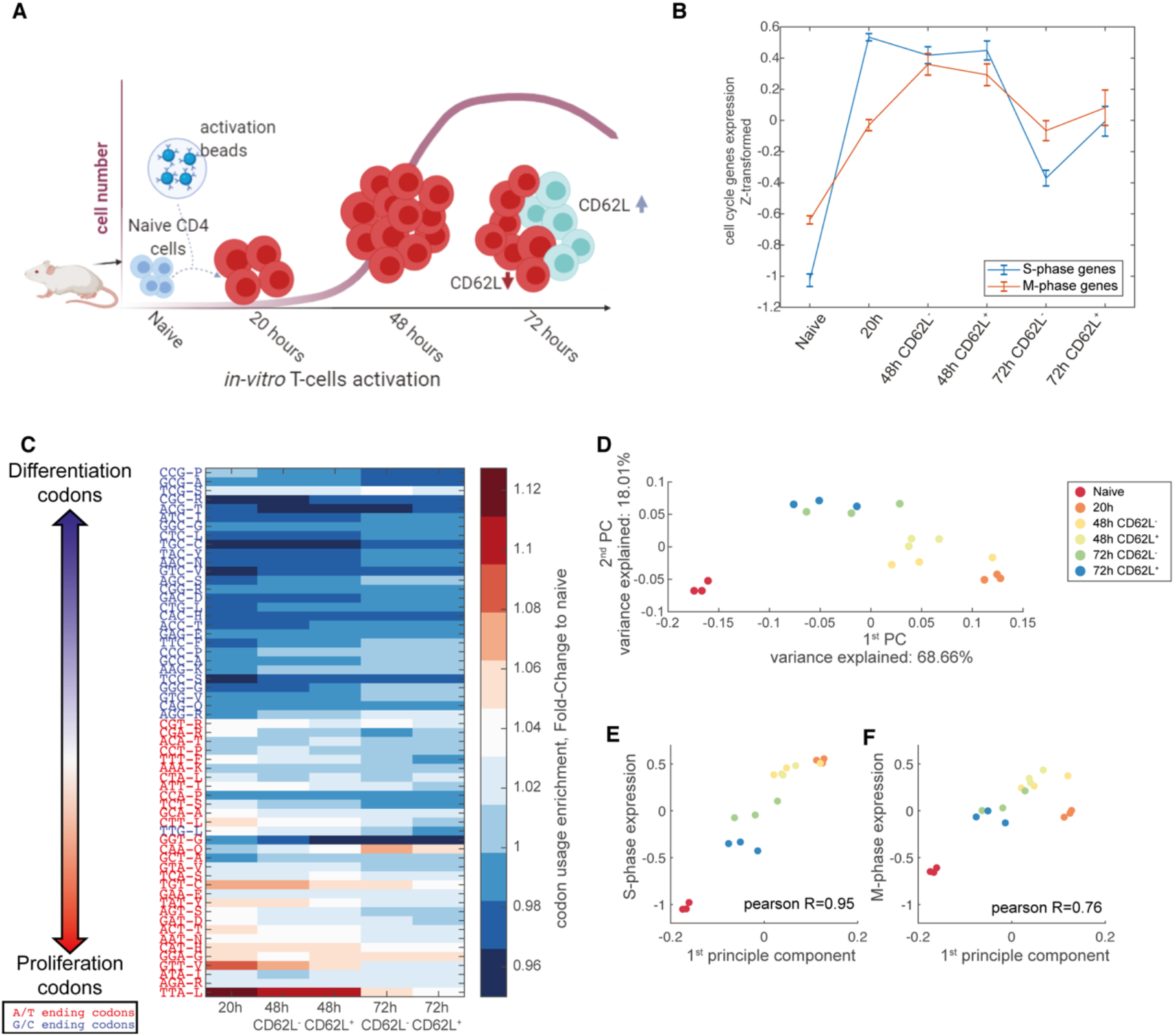
Experimental design and codon usage changes during T-cell activation. A. Schematic representation of the experimental design. CD4 T-cells were isolated from mice spleen, and activated in vitro. Naïve and activated T-cells at indicated time-points were collected and sorted for CD62L+ or CD62L-at later time points. Samples were subjected to mRNAs and tRNA sequencing. Unsorted samples were analyzed also by ribosomal protection sequencing and LC/MS shotgun analyses. B. Expression of S and M – phase genes in naïve and activated T-cells (Z-transformed average of normalized read count of S-phase pathway (147 genes) and M phase pathway (326 genes), n=3, mean ± std). C. changes in codon bias during T-cell activation process. Analysis of codon usage changes based on mRNA expression (normalized to amino-acid, i.e., “codon-bias”) as measured by RNA-sequencing. The codons are sorted based on proliferation vs. differentiation codon usage as described in Gingold et.al^5^. Codons ending with C or G are marked blue, codons ending with A or T are marked red. Sample identity is shown on the x axis (average of 3 samples, ranksum p-value <10^5^ for A/T Vs GC ending codons). D. Codon bias differentiates proliferative and arrested T-cells. Shown here is a principal component analysis of the samples, based on their codon bias, as presented in panel C. The first component (69% variance explained) best separates the naïve and 20h samples. E and F. Correlation of position in first PC plotted against average expression of S-phase (E), and M-phase (F) genes (Z-transformed, average of 129 and 292 genes respectively).

We first characterize changes in mRNA codon usage by examination of the change in representation of each of the 61 amino acid coding codons in the mRNA pool in each time point. We examined first, and verified that changes in mRNA expression were in good correlation between the two biological repeats (Fig S1-1). The “demand” for tRNA by each codon type in the transcriptome is computed as the sum, over all mRNAs, of the product of mRNA expression level and the number of appearances of each codon in each gene^5^ (see Material and Methods). We found that relative to the naïve cells, activated proliferating cells show an increase in representation of the codons defined before by Gingold et al.^5^ as the “proliferation codons” signature (Fig. 1C, Fig S1-2). The “proliferation codons” tend to end with A or T nucleotides at the third codon position, in contrast to the “differentiation codons” which tend to have G or C at their 3^rd^ positions. Although the enrichment in “proliferation codons” is seen in all samples, when compared to naïve cells, the fold change in codon usage is maximal at the 20h sample, and it is then reduced 72h after activation, as 72h CD62L^+^ cells show the least difference in codon usage from naïve cells (Fig. 1C, Fig S1-2A).

We characterized each sample based on the average codon usage of the genes expressed in it, normalized by their expression level, using Principal Component Analysis (PCA, Fig 1D Fig S1-2B). The analysis shows a marked change in codon usage at 20h, relative to the naïve population, and a gradual return towards base levels with 72h CD62L^+^ cells located closest to the naïve samples. We then turned to examine which genes derive the codon usage trend. We first checked the genes that belong to the Gene Ontology categories associated with the different stages of the cell cycle (G0-early G1, G1, S, G2M, M). We found that the first PC of the codon usage PCA is in high correlation with the expression of S-phase genes (R=0.95). The correlation to G1, G2/M, and M-phase GO categories was also high, but lower than S-phase (R=0.67, R=0.74, R=0.55, respectively, Fig. 1E,F and Fig. S1-3), while “developmental process” related genes are negatively correlated with the first PC (Fig. S1-3). Apart from cell-cycle related GO categories, the most correlative GO categories are: “negative regulation of oxidative stress-induced cell death”, “regulation of DNA metabolic process”, “cellular respiration” and “purine ribonucleoside monophosphate metabolic process” (see dataset S1). We conclude that induction of genes belonging to these functional categories drives the proliferation-codon usage signature that we observe in activated T-cells.

### Induction of a proliferation tRNA signature is followed by relaxation at the differentiation phase

We next set to examine what are the changes in the tRNA pool and if they correspond to the changes in demand as reflected by the codons usage, adapting a recently developed tRNA sequencing protocol^27^ (See Materials and Methods for further details, and Hefetz et al. for additional quality assessment). In the downstream analysis we grouped together read counts of all tRNAs of the same anticodon that have at least a minimal tRNAScan prediction score (tRNAscan-SE^28^) above 50. We calculated the expression of each such isoacceptor group as the sum of read counts mapped to each of its corresponding genes. The experiment was done in two biological independent experiments, with 3 technical repeats for each time-point, showing high correlation between repeats (Fig S2-1). Sequencing results were validated by real-time qPCR, showing similar trends for three tested tRNA types (Fig S2-2). We found a significant positive correlation between tRNA expression levels and expressed codon usage in the naïve cells, with high levels of both G/C ending codons and matched tRNA (Fig 2A, Pearson R=0.66, p-val <10^−8^). 20 hours after activation of the cells, the tRNAs that decode the A/T ending codons are upregulated compared to naïve T-cells (Fig. 2B, rank-sum test p-value<0.05). Beyond this time point, we did not observe such significant difference in expression fold change between tRNAs that code for A/T ending codons and G/C ending codons. We inspected specifically two unique tRNAs - for Seleno-cysteine and the initiator-methionine tRNA (tRNA^iMet^). We found that both these tRNAs were down-regulated at all time points following the T Cell activation (Fig 2B, Fig S2-3). While the decline of the Seleno-cysteine tRNA in proliferation is in line with previous observations, the decline in expression of the tRNA^iMet^ is contrasted with the induction of this tRNA in other proliferating cells^5,29^. We analyzed the expression of translation initiation factors and found two clusters of expression, one that is elevated upon T-cells activation and the other that is reduced, similarly to tRNA^iMet^ (Fig S2-3).

**Figure 2-.**
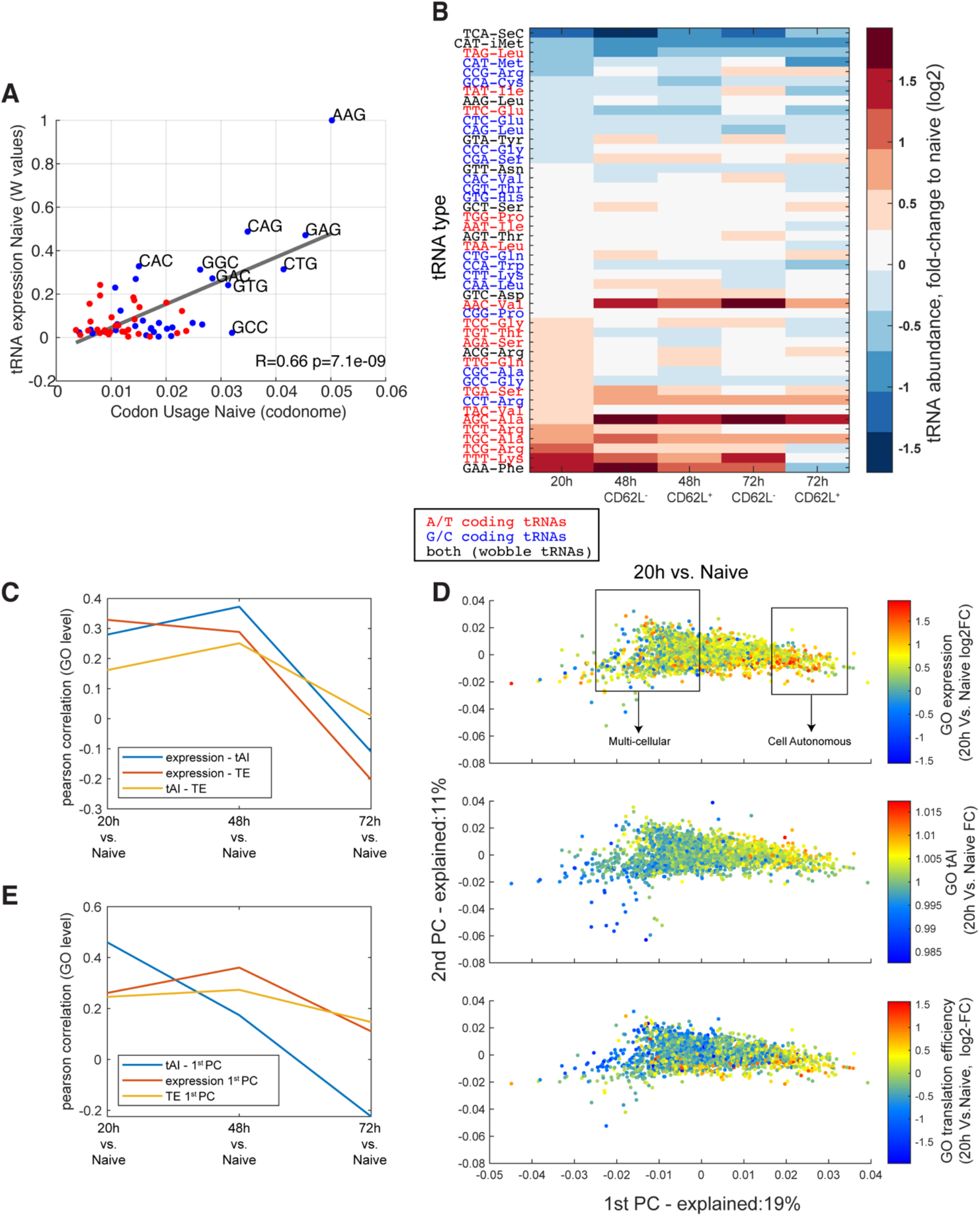
tRNA avilabilty and translation efficiency change during T-cell activation. A. Expressed codon usage (as in Figure 1C, not normalized to amino acid, x-axis) and tRNA expression (W value for codons, see Material and Methods - estimating tAI section) in naïve cells are correlated (Pearson R=0.66, p-val=9*10^−9^). Red indicates A/T ending codons, blue indicates G/C ending codons. B. Dynamics of tRNA expression during T-cell activation. Shown here is a heat-map representation of changes in tRNA expression at each time point, normalized by expression in Naïve T-cells. Rows are sorted based on the fold-change between naïve and 20h after activation. Marked in red are tRNAs that codes for A/T ending codons, marked in blue are tRNA that codes for G/C ending codons, and in black tRNAs that codes for both (by wobble interaction), iMet and Sec tRNAs. C. Pearson correlation of mRNA expression, tAI, and translation efficiency (TE) averaged for gene groups derived from GO categories (p-value for all correlations <10^−10^, except for tAI-TE at 72h (n.s.)) D. A PCA projection of the mouse codon usage of gene sets derived from GO categories. Each point represents one gene set. Gene sets corresponding to tissue-specific GO terms are to the left side, and those corresponding to proliferation related GO terms are to the right. The color code in the upper panel represent changes at the mRNA level, averaged over all the genes in each gene set. In the middle panel, each gene category is color coded according to the relative change in availability of the tRNAs that correspond to the codon usage of its constituent genes, averaged over all genes in the gene set. A red color for a given gene set indicates that on average the genes in that set have codons that mainly correspond to the tRNAs that are induced in the condition, whereas a blue color indicates that the codon usage in the set is biased toward the tRNAs that were repressed in that given condition. In the lower panel the color code indicates translation efficiency (TE) calculated based on ribo-seq by ratio of reads of the ribosome protected fragment (RPKM) divided by read count of mRNA-seq (RPKM) E. Correlation of mRNA expression, tAI, and translation efficiency (TE) to the 1^st^ PC, calculated based on average codon usage of each GO category (shown in panel C. p-value for all correlations <10^−10^).

We next aimed to reveal how coordinated are the various levels at which translation is regulated as the cells complete a cycle of proliferation and differentiation. To that end, we examined mRNA expression, tRNA expression, and translation efficiency (TE) as assessed by ribosome profiling sequencing (Ribo-seq) from the same cultures. While mRNA and tRNA seq reveal respectively the demand and supply for the translation process, ribo-seq measurement provides a snap-shot of actively translating ribosomes across the body of each translated mRNA. When analyzing the mRNA-seq and Ribo-seq libraries across coding regions, the expected distinct profiles of ribosome accumulation at the start and end of the ORF in the ribo-seq library, as well as alignment of reads to the correct ribosome reading frame, confirming the overall quality of the libraries (Fig S2-4). To assess the adaptiveness of each gene’s sequence to the tRNA supply we used a modification of the tRNA adaptation index (tAI) score^30^, which calculates codon adaptiveness based on the tRNA availability. Whereas in the original tAI measure tRNA availability is given as a static tRNA gene copy number in the genome, here, we determine the tRNA availability based on the tRNA-seq read count for each anticodon (see Materials and Methods). This allows us to determine translation efficiency for each codon in a dynamic fashion, across the process of T-cell activation. We next compute the ratio of this tAI score for each gene given the tRNA pool at a given time point, normalized to the score computed based on the naive tRNA pool. The ratio is higher than 1 at a given time point during T-cell activation for genes with a codon bias that correspond to tRNAs that are induced in that time point. tAI and TE are distinct dynamic measures of genes’ translation and the correlation between them was so far not assessed. While TE captures both initiation rate and translation speed, tAI evaluates the availability of tRNAs needed to translate a gene.

Over all, at the single gene level, we found low, yet significant correlations between changes in mRNA expression, tAI, and TE (Fig S2-5). When examining the correlation between mRNA levels and each of these measures of translation we found modest correlation (Pearson’s correlation = 0.11 but highly significant p-values due to involvement of 4474 genes in the calculation) between tAI and mRNA levels at 20h and 48h, and a slightly lower correlation between mRNA and TE at these times points. Interestingly, these correlations become (significantly) negative at the 72hr time point, perhaps suggesting the action of means to restrain mRNA’s transcription response (Fig S2-5). Yet, when analyzed at a higher level of granularity, of GO functional gene sets, we found a significant correlation between the changes in tAI, and TE at 20h and 48h after activation compared to the naïve cells (Fig 2C, Fig S2-6). Existence of such a correlation between entire functional categories but not at individual gene level suggests that different genes in shared functional categories use different regulatory means – e.g. tRNA availability or engagement with ribosome - to respond in same directions. At 72 hours after activation, there is a negative correlation between changes in mRNA expression compared to naïve and changes in TE and tAI compared to naïve.

In parallel, we examined the response of each gene and genes’ functional categories in light of their codon usage programs. For that we projected all mouse genes’ GO categories on a PCA plane, based on the average codon usage of all the genes that belong to each category (Fig. 2D). The location (but not the coloring) of the GO categories is identical in all three sub-figures, as it reflects static codon usage only. In similarity to the human genes analysis^5^, on the right hand side are cell autonomous functionality such as the cell cycle genes, and functions related to gene expression such as the ribosome, DNA replication machinery genes, while on the left side are multi-cellularity related functions (Fig S2-7). While the coefficient of the 1^st^ PC separate codons by their A/T or G/C, the 2^nd^ PC separate codons by their amino acid hydrophobicity (Fig S2-7B,C). This analysis clearly shows the marked difference in codon usage of genes that fulfil proliferation and differentiation functions, captured by the 1^st^ PCA component. To examine correspondence between changes in tRNA supply, translation efficiency and mRNA level we colored the GO categories based on averaged (Z-transformed) change in mRNA expression (Fig 2D, upper panel), based on changes in tAI calculated for averaged codon usage of the GO categories genes (Fig 2D, middle panel), and based on averaged (Z-transformed) changes in translation efficiency (Fig 2D, lower panel), between activated T-cells at 20h after activation and naïve cells. We found that the proliferation-related GO categories on the right-hand side of the PCA projection are strongly induced at the mRNA level at 20h after activation compared to naïve cells. The signature is stronger at 48 hours, and reduced at 72 hours after activation, reflecting reduction in proliferation gene expression at 72 hours (Fig 2E, Fig S2-8). The dynamics of changes in translation efficiency of GO categories, in correlation with the 1st PC component, recapitulates the changes in mRNA expression (Fig 2E, Fig S2-8). The correlation of changes in tAI with the 1^st^ PC calculated for GO categories is highest at 20h after activation compared to naïve, and become negative at 72h after activation (Fig 2E, Fig S2-8). Hence, at 72 hours after activation, the proliferation-related genes are still induced at the mRNA level, but to a lesser extent compared to the 20 hours, and the tRNA pool supply is no longer biased towards the codon usage of the genes that belong to these functionalities. This is accompanied with negative correlation between changes in mRNA expression to translation efficiency and tAI (r=-0.1, r= -0.2, respectively, Fig 2C). The relationships between the tRNA and mRNA pool at 20 hours, but not in 72 hours, in which genes that serve cell autonomous functions are induced at the mRNA level, and tRNAs whose anticodons match their codon usage are induced as well, resemble the situation seen in diverse cancer types^5^. It is possible to speculate that the lack of correlation between supply and demand at the 72 hours time-point might ensure that the pro-proliferation mRNAs that may still exist in the cell at this stage will not be translated efficiently, restraining potential undesired excessive proliferation. In summary, this analysis reveals some level of coordination between mRNA and tRNA supply and demand during the dynamic process of T-cell activation, yet this response relaxes, and with it this coordination, and it also shows that functionally related genes might use different regulatory means to respond in similar directions.

### tRNA modifications involved in translation frame maintenance are reduced in activated T - cells

tRNA molecules are extremely rich with post-transcriptional RNA modifications, of which some can govern the stability, translation efficiency, decoding rate and fidelity^11,14,31,32^. Chemical modifications of RNAs are reliably detected and quantified by means such as mass-spectrometry^33^, and in particular mass-spectrometry has been used to analyze tRNAs 9,22,34. An interesting question is can modifications on RNAs be detected using RNA-seq methods, given that many modifications can often cause failure in the reverse transcription (RT) reaction^13^? In modified positions, the reverse transcriptase can either transcribe the original nucleotide, abort the reaction, resulting in truncated fragments or it may introduce typical nucleotide mismatches compared to the genomic sequence. Indeed in several protocols for sequencing of RNA, and tRNAs in particular, certain modifications are enzymatically removed from the transcription prior to sequencing^35,36^. Yet, in agreement with recent works^13,37^, we realized that beyond our ability to uniquely detect and quantify a modified tRNA, the disadvantage due to RT disruption can actually turn into an opportunity – to quantify the modification level.

To calculate the percentage of modified tRNA, as these may be dynamically regulated, we relay of the fact that the fraction of RNA copies that carry the modification can be calculated as the sum of mismatches and RT–abortion truncated reads at the position, normalized to the total number of reads that map, either perfectly or not, to the gene sequence. Modifications that do not interfere with the RT process are not detectable through this method. We note that the data that emerge from sequencing do not allow to fully detect the actual chemical nature of the modified nucleotide, only relative changes in the level of editing, but crossing the read data with databases of annotated modifications may allow to infer the nature of modification and to detect changes in the fraction of edited transcripts. We note that our estimation of modification levels, that is based here on percentage of truncated reads, and reads with mismatched nucleotides, provides only a lower bound on the modification level since it is possible that some of the modified RNA positions will be reversed transcribed with no errors or abortions.

To determine which modifications can be detected using tRNA-sequencing we crossed annotated modification positions from the MODOMICS database^9^, with tRNA-sequencing data collected in this study and human^8^ and *E. coli*^38^ data from previous studies. We found that out of 46 modification types, annotated in MODOMICS for the indicated species, 13 modifications are detected based on mismatches or RT-abortions in our protocol (table S1, Figure S3-1). Undetectable annotated modifications are either not causing mismatches or RT-abortions, or they may have not been sufficiently modified in the sample measured. For mouse, only twelve tRNA species are annotated with their modifications in the MODOMICS database, and in total they represent 18 chemical modification types, appearing in 196 modification positions in total. Through changes in tRNA reads we have been able to detect 5 out of 18 annotated modifications (m1A, m1G, m2,2G, I, and wybutosine), in 24 known positions. In addition, 615 un-annotated sites appear to be at least minimally modified (at least 10% of the reads, see detailed modification list – dataset S2).

Based on modification calling by mapping mismatches and RT-abortion at each position, we could quantify changes in modification levels throughout the T-cell activation process. We examine separately the modifications around the anticodon (and one nucleotide upstream and downstream), which are known to have a higher effect on tRNA decoding role^39^ (Fig 3A), and the modifications that occur elsewhere in the tRNA molecule, which are involved in addition in many other aspects of tRNA biology^40^ (Fig. S3-2). We found that the level of most modifications remained constant during T-cell activation process. However, two types of modifications located on tRNA^Lys^(UUU) and tRNA^Phe^(GAA) show marked reduction in modification level in activated cells at 20 hours compared to the naïve cells. The two modifications are within the anti-codon loop, on position 37, i.e. one nucleotide upstream from the anticodon, (Fig. 3A). We also detected changes in modification levels of tRNA^Arg^(UCU) isodecoders at a position more distant from the anti-codon. According to the annotation in MODOMICS this modification can be assumed to be m^3^C 32^13^ (Fig. S3-2). We focus below on the modifications in tRNA^Lys^(UUU) and tRNA^Phe^(GAA).

**Figure 3-.**
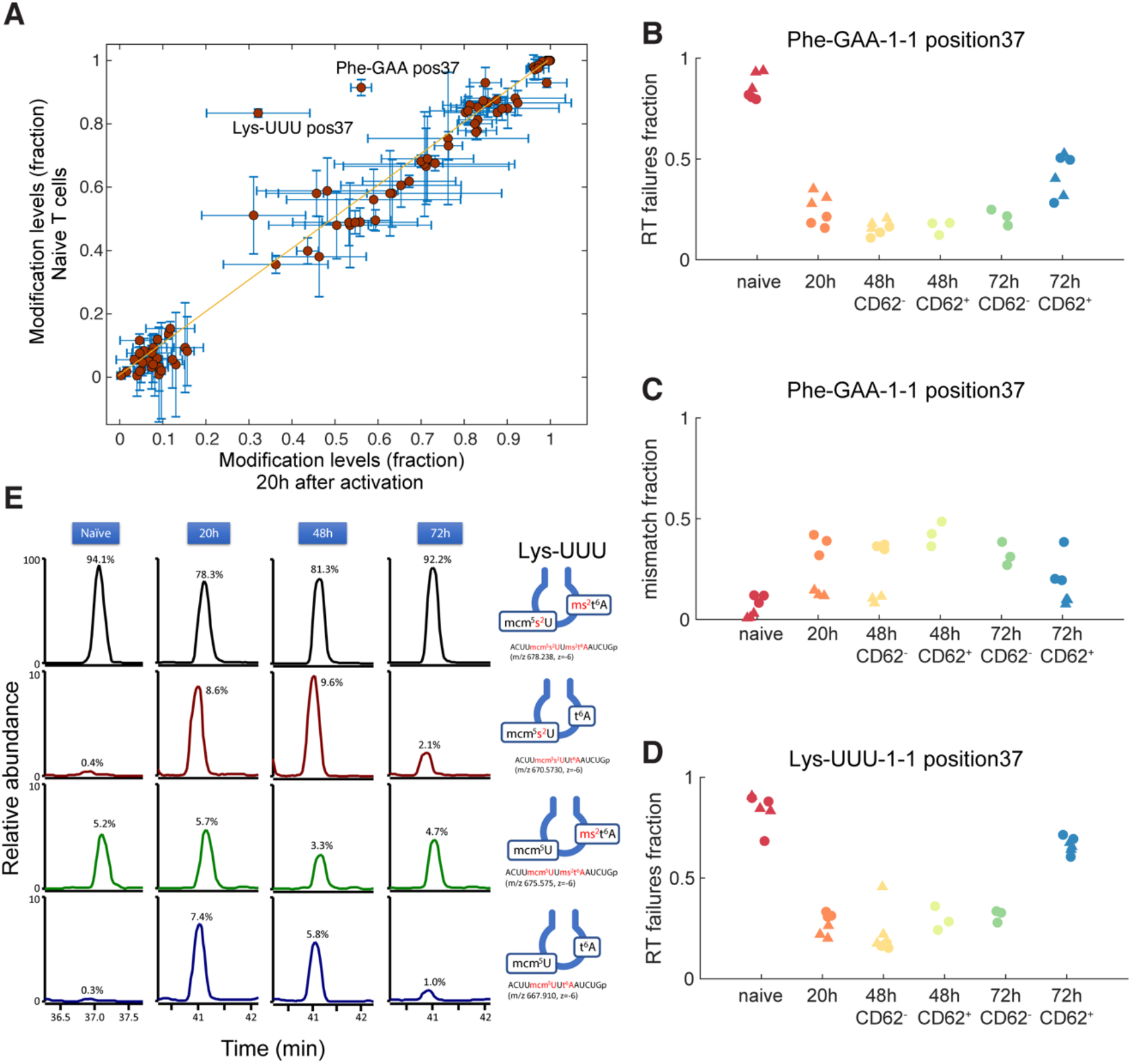
Changes in tRNA modifications during T-cell activation. A. Average anticodon loop modification levels based on tRNA-seq from the naïve and activated (20h) T-cells (n=3 ±standard deviation). Threshold for modification calling is: minimum coverage of modified position = 20 reads, modification fraction above 10% in at least one sample. B. Reverse transcription-abortion fraction at position 37 of reads aligned to tRNA^Phe^(GAA). Triangles and circles are the first and second experimental sets correspondingly. (anova p-value <10^−5^ for all samples compare to naïve, <0.01 for all samples except for naïve compared to 72h CD62^+^). C. Mismatches fraction at position 37 of reads aligned to tRNA^Phe^(GAA) (anova test for first set: p-value <10^−4^ for all samples except for 72h CD62^+^ compare to naïve, naïve vs. 72h CD62^+^ p-value<0.05, second set-n.s). D. Reverse transcription - abortion fraction at position 37 of reads aligned to tRNA-LysUUU. (anova p-value <10^−7^ for all samples except for 72h CD62^+^ compare to naïve, naïve vs. 72h CD62^+^ n.s, p-value<10^−7^ for all samples except for naïve compared to 72h CD62^+^). At this position, there is no detected changes in mismatches levels (Fig. S3-3). E. Mass spectrometric shotgun analysis of cytoplasmic tRNA modifications during T-cell activation. Shotgun analysis of class I tRNA fraction from mouse CD4+ T-cells collected at 0 (naive), 20, 48 and 72h after activation. The tRNA fraction was digested by RNase T1 and subjected to capillary LC/MS. Top to bottom panels represent XICs for negative ions of anticodon-containing fragments of cytoplasmic tRNA^Lys^ bearing mcm^5^s^2^U34 and ms^2^t^6^A37, mcm^5^s^2^U34 and t^6^A37, mcm^5^U34 and ms^2^t^6^A37 and mcm^5^U34 and t^6^A37, respectively. The percentage of each peak represents relative abundance of the corresponding fragment.

Interestingly, both of the anticodon modifications that changed in level during T-cells activation are involved in decoding of “slippery codons” which are known to be involved in ribosomal frameshift^17,41–43^. Position 37 of tRNA^Phe^(GAA) is annotated in MODOMICS to have the wybutosine modification. wybutosine (yW) and its derivatives, peroxywybutosine (o2yW) and hydroxywybutosine (OHyW), are tricyclic nucleoside with a large side chain found at position 37 of mammalian tRNA^Phe^. yW derivatives play a critical role in efficient codon recognition and reading frame maintenance^17,43^. We found that RT-abortions fraction at position 37 of tRNA^Phe^(GAA), assumed to result from the wybutosine or derivates presence, changed dynamically with T-cells activation. Its levels were high at naïve T-cells, low at 20h and 48h after activation, and elevated back at 72h after activation yet to a lower level compare to initial level (Fig 3B). While RT-abortion due to the modification decreased during T-cell activation, mismatches fraction increased in the activated cells (Fig. 3C), showing a distinct mismatch pattern (Fig S3-3A) which is similar to the pattern observed in annotated positions of m1G modifications (Fig S3-1). This indicates that in the activated cells m1G modification is not processed to wybutosine. However, we did not observe any significant reduction in mRNA expression of wybutosine biosynthesis enzymes (tyw-1, trmt-5, tyw-3, lcmt-2, tyw-5, dataset S3). For tRNA^Lys^(UUU) we found RT-abortions in a similar pattern to tRNA^Phe(^GAA), without the appearance of mismatches in the activated cells (Fig 3D, Fig S3-3B). The intermediate modification t^6^A, that might occur in the activated cells, cannot be detected using our tRNA-sequencing protocol, since it does not result in a detectable mismatch pattern (Fig S3-1). Our data support the presence of the bulky modification 2-methylthio-6-threonylcarbamoyl-A (ms^2^t^6^A) as shown before in mouse^18^ and human^9,44^, as this modification was shown to cause RT abortion in human^13^. while the two above mentioned modification (i.e t yW on tRNA^Phe^(GAA) and ms^2^t^6^A on tRNA^Lys^(UUU)) show dynamic change in levels during T-cell activation, the downstream modification m1A at position 58 remained constant for both tRNAs (Fig. S3-3C). The retainment of this modification on both tRNAs indicates that these tRNAs maturate and are processed properly.

We were looking for a sequencing independent means to identify and quantify these modifications and turned to the direct analysis of tRNA modifications by mass spectrometry. For that, class I tRNA fractions were obtained from the naïve CD4+ T-cells and activated cells at 20h, 48h and 72h. Each tRNA fraction was digested by RNase T1 and subjected to the shotgun analyses by capillary liquid chromatography/mass spectrometry (LC/MS)^45,46^. As shown in Figure 3B, we successfully detected the anticodon-containing fragments of cytoplasmic tRNA^Lys^(UUU). Judging from the exact molecular mass, we found four kinds of fragments with different modification status. The fully modified fragment containing 5-methoxycarbonylmethyl-2-thiouridine (mcm^5^s^2^U) at position 34 and 2-methylthio-*N*^6^-threonylcarbamoyladenosine (ms^2^t^6^A) at position 37 was detected as a major species in all four time points during the activation process (Fig. 3E). This fragment was further probed by collision-induced dissociation (CID) to confirm the position of each modification (Fig. S3-4). In addition, we detected three hypomodified fragments bearing mcm^5^s^2^U34 and t^6^A37, mcm^5^U34 and ms^2^t^6^A37, and mcm^5^U34 and t^6^A37 (Fig. 3E). Over time after activation, these hypomodified fragments increased significantly, suggesting that 2-methylthio group of ms^2^t^6^A37 and 2-thio group of mcm^5^s^2^U34 are down-regulated at 20h and 48h after activation. Since 2-methylthio group of ms^2^t^6^A37 hinders reverse transcription and can be detected by RT stop signature, The LC/MS data clearly explain the unique behavior in the tRNA-sequencing data upon T-cell activation.

In the shotgun analysis of CD4+ T-cells, we clearly detected the anticodon-containing fragments of tRNA^Phe^. OHyW is a major modification and o2yW is a minor modification in naïve cell (Figure S3-5). Upon activation, both yW derivatives get plunged drastically (Figure S3-5), indicating that biogenesis of yW derivatives is impaired after T-cell activation. This result nicely explains the deep sequencing data for tRNA^Phe^(GAA) upon T-cell activation.

### Reduced wybutosine modification level increases ribosomal frameshift level

Since we could not detect intermediate modifications of yW derivatives from tRNA^Phe^ by mass spectrometry, and to estimate the effect of the modification on ribosomal frameshift, we first approached this modification by a combined tRNA sequencing and a gene deletion approach. Specifically, we used CRISPR-cas9 to knock out a wybutosine modification enzyme, the gene *tyw1*. This enzyme catalyzes the second reaction in the biogenesis of yW formation at position 37 of tRNA^Phe^. We generated clones of HeLa *tyw1* knock-out-cells using two alternative gRNAs that target the second or third exon of *twy1* gene. Indeed, the tRNA sequencing-based signal for detection and quantification of wybutosine modification level was reduced on tRNA^Phe^(GAA) by 20 to 30%, from an initial level higher than >90% (Fig. 4A). Interestingly, the total level of tRNA^Phe^(GAA) is elevated upon this knock-out (Fig. 4B), suggesting a feed-back loop that could sense the reduction in the modified tRNA level and respond by elevating transcription of the tRNA molecules. Alternatively, it is possible that the modification de-stabilizes the tRNA.

**Figure 4-.**
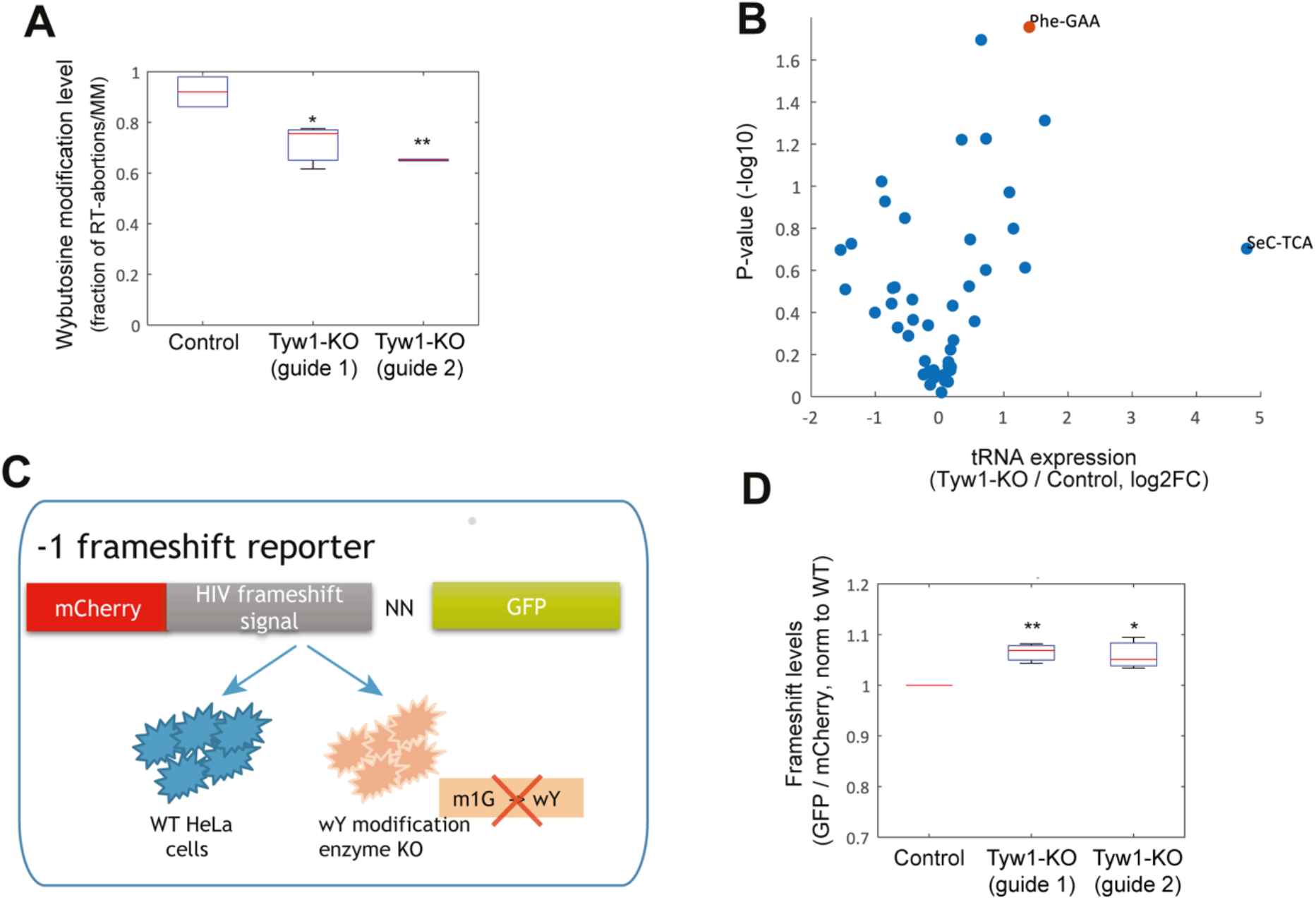
Reduction of wybutosine modification induces translation frameshift. A. Changes in modification levels, measured by tRNA sequencing. Shown are fraction of RT abortion at position 37 of reads aligned to tRNA^Phe^(GAA) in control cells (non-targeting gRNA) as well as levels measured for two clones of knockout for tyw1 gene. Asterisks mark the statistical significance between control levels and KO, student t-test * P<0.05, ** p<0.005 (n=3 biological repeats) B. Changes in tRNA abundance. A volcano plot showing the log2 fold change of abundance levels of each tRNA type measured in WT vs. *tyw-*1 knockout. tRNA abundance is measured by the number of reads aligned to each tRNA, grouped by anticodon. Mark in red is tRNA^Phe^(GAA). C. A scheme for creating HeLa cells Knockout and measuring frameshift using a HIV reporter. D. Frameshift levels as measured by GFP expression normalized to mCherry. Shown are frameshift levels in control cells (non-targeting gRNA) as well as levels measured for two clones knocked out for tyw1 gene. Asterisks mark the statistical significance between control levels and KO, student t-test *P<0.05, ** p<0.005 (n=3 biological repeats).

Here too we turned into LC/MS – based chemical confirmation of the modification. We performed a shotgun analysis of tRNA fraction obtained from HeLa cell, and detected anticodon-containing fragments bearing o2yW and OHyW from cytoplasmic tRNA^Phe^ (Figure S4-1). The peak intensities estimate that 90% of tRNA^Phe^ has OHyW, and the rest 10% has o2yW. Any other intermediates during yW biogenesis are not detected (Figure S4-1). When *tyw1* was knocked out, both yW derivatives disappeared (Figure S4-1). Because TYW1 is responsible for conversion from m^1^G to imG-14 in yW biogenesis^45^, we expected to detect m^1^G-containing fragment (ACmUGmAAm^1^GAUCUAAAGp). However, we failed to detect it unexpectedly (Figure S4-1). As m^1^G can be digested by RNase T1, we next searched for the corresponding fragments ACmUGmAAm^1^Gp and AUCUAAAGp, and only found AUCUAAAGp whose sequence was confirmed by CID (Figure S4-2), indicating that biogenesis of yW derivatives is impaired at the second step catalyzed by TYW1 in the *tyw1* KO cell.

To test the influence of the wybutosine modification on frameshift level we used a fluorescent protein-based frameshift reporter (see Material and Methods, Fig 4C and Fig S4-3). We have recently created a library of such dual fluorescent proteins connected with diverse candidate linkers^47^. To select from this library one biologically meaningful linker sequence we returned to the biology of activated T-cells. We recall that infectivity of HIV-1 is primarily focused on activated T-cells, and is heavily dependent on the UUU (-1) frameshift for translation of its polymerase enzyme^25^’^48^. We have thus chosen the HIV gag-pol frameshift sequence as the linker for our reporter (Fig. 4C).

We transfected the *tyw1* KO cells as well as control cells, with the fluorescent frameshift reporter and measured the frameshift level. We found that the frameshift level is significantly elevated by 6% in the mutant compared to the wild-type (Fig. 4D). This indicates that down-regulation of wybutosine modification can indeed exert an effect on frameshift in general and that it might affect production of HIV polymerase, which depends on ribosomal frameshift.

### A Trade-off between translation fidelity and speed

Why do activated T-cells reduce the level of the two tRNA modifications that have a role in reducing ribosomal frameshift? We hypothesize that the T-cell system may have evolved to trade-off translation fidelity and speed, as discussed in other contexts^49,50^. One possibility is that this reduction in modifications allow cells to translate proteins more rapidly, a necessity under conditions that require rapid growth in biomass production, towards cellular proliferation. Under such conditions, we reason, the system evolved to compromise translation fidelity, i.e. to feature enhanced frameshift. This hypothesis raises the prediction that under such a trade-off scenario, genes involved in T-cell activation would avoid the slippery codons (UUU) in favor of synonymous codons (UUC) that are not prone to frameshifting, thus reducing translation error rate. We examined each mouse gene for its tendency to prefer either UUU or UUC for encoding Phenylalanine, and ranked all genes according to this codon bias. Strikingly, we found that the genes with the highest UUC/UUU ratio are enriched with several functionalities involved in T-cell and other immune functions (Fig. 5). As a negative control, we examined the other “2-box” codon duplets in the genetic code, i.e. all pairs of synonymous codons of the type XXU XXC. For each such pair, we computed across all genes, the XXU/XXC preference score and ranked the genes according to the extent of XXU avoidance. None of the other eight codon pairs showed an enrichment for any functionality related to T-Cells or immunity (table S2). Avoidance of frameshift-prone codons in the genes expressed in proliferating T-cells may reduce undesired frameshifting events that can occur due to reduction in the tRNA modification.

**Figure 5-.**
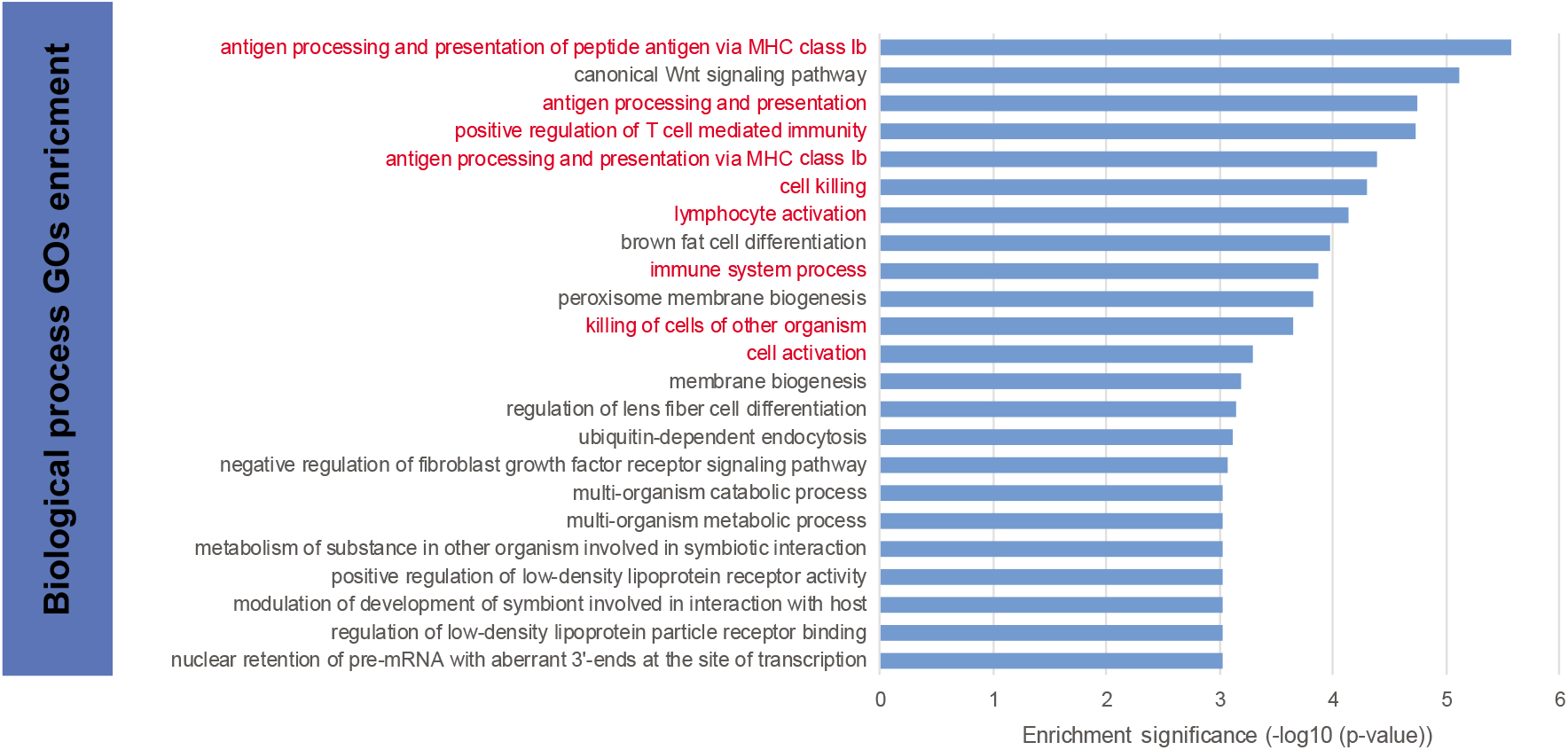
Trade-off between translation speed and accuracy. T-cell activation and immune-related genes are avoiding the use of UUU codons. Shown here are GO categories enriched with genes that avoid the use of UUU codon normalized to Phenylalanine usage. Analysis of other “U” ending codons did not show enrichment for T-cell related GO categories (table S2)

## Discussion

In this work we studied the tRNA pool and its interaction with the mRNA pool in a natural physiological context of massive cellular proliferation, that of activated T-cells. The interplay between the tRNA and mRNA pool in cells was shown to govern proteome-wide translation ^7,51,52^. Previous studies have shown that the mammalian proliferation state affects the cellular tRNA pool, but most studies were focused on cancerous proliferation^15,53^. Non-cancerous, physiologically normal proliferation may obey different dynamics. Normal cellular proliferation must be a restrained process, one that actually must have evolved to avoid undesired cancerous transformation. Normal proliferation evolved over long organismal evolutionary time scales, as opposed to cancer, which operates selection over short time scales of cellular somatic evolution. The motivation to inspect the tRNA pool under conditions of normal proliferation is thus clear.

Indeed, we find that early on upon stimulation, proliferating T-cells feature transcription activation of the proliferation translation program. Induced mRNA coding genes that fulfil the proliferation needs of cells represent a proliferation codon usage program and they are rather adequately served by the proliferation-oriented tRNAs that are largely induced at this stage. Yet, after the pick of proliferation (72hr in this experiment), while many of the proliferating mRNA transcripts are still present, the tRNA pool appears to have relaxed back and it might no longer translate as efficiently the pro-proliferation transcripts. At this time-point the difference between the CD62L^+^ and CD62L^-^ T-cells is interesting, while the tRNA pool of CD62L^+^ cells appears almost fully relaxed, the CD62L^-^ still show some residual induction of the proliferation tRNA pool signature. This is in line with reduced proliferation rate that was observed for memory precursors compared with effector T cells during influenza infection^54^. Thus, the faster decline in expansion of memory T cells might be regulated in part by dynamics of tRNA expression that result in imbalance between codon supply and demand.

This study goes beyond changes in tRNA levels, as it inspects dynamics in tRNA modifications during the T-cell activation process. While most modifications on most tRNAs do not change, we found two modification that are one nucleotide downstream of the anti-codon of two tRNAs, which drop significantly during the pick of proliferation, and are relaxed back towards base level when cells differentiate into memory cells. Interestingly, these two tRNAs decode “slippery codons” that are prone for ribosomal frameshifting^17,41,43^. It thus appears that during the pick of proliferation, cells trade off translation adequacy, and maybe even run into the risk of ribosomal frameshifting.

Why do activated T-cells reduce the level of the two tRNA modifications that have a role in reducing ribosomal frameshift? One possibility is a putative beneficial effect of frameshifting on these cells at the proliferation stage. It is possible to speculate that ribosomal frameshifting might enhance phenotypic variability of immune cells. Yet, our examination of codon usage in immune related genes (Fig. 5) does not provide further support to this notion. A more conservative alternative is that the T-cell system may have evolved to balance the trade-off between translation fidelity and speed, as has been discussed in other cellular contexts^49,50^. According to this possibility, reduction in modification allows cells to translate proteins more rapidly, i.e. the presence of the modifications stabilize the codon-anticodon pairing and increases fidelity^18,55–57^. This is a necessity under conditions that require rapid growth in biomass, towards extensive cellular proliferation. In support of this possibility, we indeed found that genes involved in immune response, such as TCF7, CD5, Lyl1, Pdcd1, and Rhoh, tend to avoid the use of the slippery codon UUU. Notwithstanding, despite the apparent avoidance of slippery codons in the immunity-related genes, other genes expressed in T-cells might still be subject to elevated frameshifting upon reduction in the modification. Thus, the second prediction of the speed-fidelity tradeoff model is that if indeed T-cells experience a general, perhaps non-specific, increase in frameshifting, they could suffer from a proteotoxic stress. Interestingly we found that the proteasome, both the catalytic and regulatory sub-units, is induced around 20h upon activation, followed by relaxation at later time points, suggesting that a proteotoxic stress is experienced by the proliferating cells (Fig S1-3, bottom panel) ^58,59^. The model that thus emerges is that T-cells reduce the level of two tRNA modifications that when present prevent frameshifting at slippery codons. In doing so, they might trade-off translation fidelity and speed. They protect their most crucial and cell specific genes from containing the slippery codons, but they may suffer from otherwise non-gene specific frameshifting activity.

Most observations in this study were done in mouse T-cells. Yet, assuming the T-cell proliferation in human cells features similar reduction of the same tRNA modifications, would lead to the hypothesis that the increase in the potential for ribosomal frameshifting may also occur in proliferating human T-cells. Interestingly, activated human T-cells are the infection targets of HIV, which necessitates calibrated frameshifting rates on the UUU slippery codon between its gag and pol ORFs^25^. Thus, while T-cells must be reducing tRNA modifications for their own needs and interest, it is tempting to speculate that HIV takes advantage of this dynamics that is utilized by the virus as a vulnerability.

## Material and methods

### T-Cell isolation and FACS sorting

Two independent biological repeats were done as following. Naïve cells were extracted from 6 and 21 spleens of B6 mice using the StemCell CD62L+CD44+ kit (female, 7 weeks, 6-9% naive). Each two (out of the 6) or seven (out of the 21) spleens were pooled to a total of 3 samples extracted separately. Naïve cells were activated by addition of anti-CD3/anti-CD28 activation bead (1:1 bead:cell ratio, thermoFisher Scientific Cat#11131D). Cells were grown at 0.25 cell/ml in 24 wells, and were collected at 20h, 48h, and 72h after activation.

For the first biological repeat, a portion from the naïve state was taken to FACS sorting for live cells followed by RNA extraction. At 20h only live cells were sorted. At 48 and 72 h cells were sorted into CD62L+CD44+ and CD62L-CD44+, as described in SI appendix, Extended Material and Methods.

### mRNA sequencing

mRNA sequencing for the first biological repeat was preform using MARS-Seq protocol as described ^60^ and analyzed as described in SI appendix, Extended Material and Methods. Libraries were sequenced using 75bp single read output run on illumina NEXTseq platform.

mRNA-seq for the second biological repeat was done in parallel to ribosome footprint sequencing by a tailored mRNA-sequencing protocol as previously described ^61^(see and SI appendix, Extended Material and Methods for further details

### tRNA sequencing and read analysis

tRNA sequencing protocol was adapted from Zheng *et al*.^36^. with minor modification, described in SI appendix, Extended Material and Methods. The protocol involves the use of a highly persistent and thermo-stable reverse transcriptase. The primers are DNA-RNA hybrids that have overhang of thymidine, which enable reverse transcription of adenosine ending RNAs. This leads to enrichment of mature tRNAs, due to their shared CCA tail. Using this tRNA sequencing protocol, 45%-60% (depending on the sample) out of the reads are mapped to mature tRNAs, the cytosolic or mitochondrial. Additional 1%-6% of the reads were aligned to pre-mature tRNAs. Out of 273 unique mature tRNA sequences in the mouse genome, 120-156 tRNA genes get sufficiently high read counts that allow their quantification. On the other hand, 116-141 tRNAs did not get sufficient number of reads (under 20 reads). All the tRNA genes with a low scores (below 50, tRNA-Scan SE^28^), have low read counts, and can be considered as pseudo-tRNA (Fig. S5-1A). The low read count indicates that those tRNAs are either not expressed or cannot pass through the maturation process. Individual tRNAs were next grouped by anticodon. We calculated the gene copy number for each anticodon based on the number of tRNA genes in the genome, with prediction score above 50. We found that although they exist in the genome, tRNAs of certain anti-codons, with low gene copy number, have very low expression (Fig. S5-1B). For all of those anticodons, the matching codons can be translated by wobble interactions with a different tRNA group. These results are similar to measurements done in human HEK293 cells and in the brain^62^, and they are in agreement with the predication that approximately a third of the tRNA genes in mouse are inactive in all tissues, and around 30% are tissue specific^63^.

### Codon usage calculation

Codon usage and codon bias were calculated based on mRNA expression. We summed the usage of each codon in each gene multiplied by the expression of the gene.

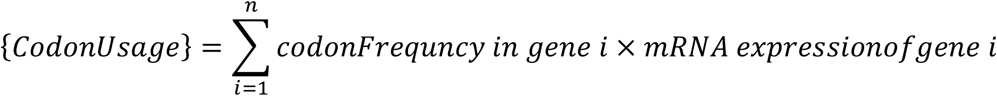

### Estimating tAI in based on tRNA expression

We generated a new measure of translational efficiency (implemented in Figure 2). Our measure is calculated similarly to the tAI measure of translation efficiency^30^, with one major change-we determine tRNA availability based on the read aligned to the indicated tRNA, as done in our previous work5. As such, the new measure can be computed for every condition. we defined the tRNA availability of each tRNA type (anticodon) by the sum reads of its tRNAs. Then, we determined the translation efficiency of an individual codon by the extent of reads of tRNAs that serve in translating it, incorporating both the fully matched tRNA as well as tRNAs that contribute to translation through wobble rules^64^ (W-value).

Since there are biases in the tRNA sequencing methods we employed this method as a comparative approach-we report fold-changes between conditions and not the actual value.

### Frameshift assay

Control and TYW-1 knock-out HeLa cells were seeded onto 6-well plates such that cell confluence will be approximately 70% the next day (∼150,000 cells). Cells were transiently transfected with frameshift reporter plasmids. The plasmids express mCherry, followed by HIV gag-pol frameshift signal, followed by GFP out of frame (-1). As a control we used in-frame GFP and out of frame (+1) GFP in the same plasmid context (Fig 4C and S4-3A). The plasmids were generous gift from Martin Mikl^47^. The cells were analyzed using ATTUNE FACS (ThermoFisher) 48h after transfection. Frameshift rate was calculated as GFP/mCherry ratio, in the linear range of fluorescent slop (Fig. S4-3).

SI Appendix, Extended Materials and Methods include further details of the study materials and methods.

### Data availability

Sequencing data are available in GEO under the accession code GSE165622.

### Data sources

#### Coding Sequences

The coding sequences of M.musculus were downloaded from the Consensus CDS (CCDS) project (ftp://ftp.ncbi.nlm.nih.gov/pub/CCDS/).

#### Classification of Gene Categories

Defined gene categories by biological process were downloaded from the Gene Ontology project (http://www.geneontology.org/); to avoid too-small gene sets, we only considered those with at least 40 genes. Pathway lists were downloaded from mouseMine.

#### tRNA modifications

Annotation of tRNA modifications were download from MODOMICS^9^ database (http://modomics.genesilico.pl)

## Acknowledgements

We are thankful to the European Research Council, the Israel Science Foundation and Israel Cancer Research Fund for grant support. We thank the Pilpel and Friedman lab members, Noa Hefetz-Aharon, Martin Mikl, Schraga Schwartz, and Daniel Douek for stimulating discussions

