## Supplemental Information for "Dynamic changes in tRNA modifications and abundance during T-cell activation"

#### Classification:

BIOLOGICAL SCIENCES; cell biology

#### This PDF file includes:

- Extended material and methods
- 19 supplementary figures
- 2 supplementary tables
- 3 Datasets Legends

### Extended Material and methods

#### T-Cell isolation and FACS sorting

Two independent biological repeats were done as following. Naïve cells were extracted from 6 and 21 spleens of B6 mice using the StemCell CD62L+CD44+ kit (female, 7 weeks, 6-9% naive). Each two (out of the 6) or seven (out of the 21) spleens were pooled to a total of 3 samples extracted separately. Naïve cells were activated by addition of anti-CD3/anti-CD28 activation bead (1:1 bead:cell ratio, thermoFisher Scientific Cat#11131D). Cells were grown at 0.25 cell/ml in 24 wells, and were collected at 20h, 48h, and 72h after activation.

For the first biological repeat, a portion from the naïve state was taken to FACS sorting for live cells followed by RNA extraction. At 20h only live cells were sorted. At 48 and 72 h cells were sorted into CD62L+CD44+ and CD62L-CD44+. The cells were stained as followed and analyzed using ARIAIII FACS:

| **Antibody** | **fluorophore** |
|---|---|
| CD4 | Pacific-Blue |
| CD62L | PE/Cy7 |
| CD44 | FITC |
| CD69 | PE |
| CD25 | APC |

#### RNA extraction

Sorted cell and unsorted culture cells were washed with PBS twice. The supernatant was removed and cells were re-suspended in TRIzol (thermofisher scientific). RNA was extracted according to manufacture protocol.

#### Cell culture

HeLa cells (ATCC; CRL-3216) were grown in DMEM high glucose medium (Biological Industries; 01-052-1A) supplemented with 10% FBS, 1% penicillin/ streptomycin (P/S) and 1% L-Glutamine. Transfections were done using JetPEI transfection reagent (polyplus transfection) using standard manufacture protocol.

#### Generation of *tyw1* knockout

HeLa cells were seeded onto 10cm plates and grown for 24h until reaching a confluence of approximately 70% (seeding ~$2*10^6$ cells). 5μg of LentiCRISPR V2 vector (addgene #52961) containing either one of two sgRNA for *tyw1* gene or non-targeting control sgRNA was transfected to cells using JetPei transfection reagent. Guide RNA sequences was chosen from Predesigned Alt-R® (IDT).

Guide sequences are:

**TWT1 guide 1**: 5'-AACCACTCTGCACTTTCAGT,-3'

**TWT1 guide 2**: 5'-TCTTCACTGGAGCTCTCAAA-3'.

**Non-targeting guide**: 5'- TTTCGTGCCGATGTAACAC-3'.

Four hours after transfection, the medium was replaced with a fresh medium. Two days after the transfection, medium was replaced with selection medium (DMEM containing 2 μg/ml puromycine) and refreshed every day for 3 days until non-transfected cells reach 100% death (no adherent cells remained in dish). Transfected cells were diluted in 96 wells plate to a concentration of a single cell per well and grown for two weeks with selection medium to generate single clones. Knock-out of *tyw1* gene in each clone was validated using sanger sequencing with the following primers:

For *tyw1*- guide 1 validation:

Fwr: 5'-GGTGCCCTTTGTTCTTTGAGATG-3'

Rev: 5'-GTGTGCGTGTGTGCACGTG-3'

For *tyw1*- guide 2 validation:

Fwr: 5'- GAGAGCTTTGCCTTTGCCCC -3'

Rev: 5'- CTTTTCTTTCTTCACATGATCC -3'

We arbitrary selected clones that show editing in the PAM position (Fig S4-3B) for downstream analysis.

#### Frameshift assay

Control and TYW-1 knock-out HeLa cells were seeded onto 6-well plates such that cell confluence will be approximately 70% the next day (~150,000 cells). Cells were transiently transfected with frameshift reporter plasmids. The plasmids express mCherry, followed by HIV gag-pol frameshift signal, followed by GFP out of frame (-1). As a control we used in-frame GFP and out of frame (+1) GFP in the same plasmid context (Fig 4C and S4-3A). The plasmids were generous gift from Martin Mikl[1]. The cells were analyzed using ATTUNE FACS 48h after transfection. Frameshift rate was calculated as GFP/mCherry ratio, in the linear range of fluorescent slop (Fig. S4-3).

#### mRNA sequencing

mRNA sequencing for the first biological repeat was preform using MARS-Seq protocol as described [2]. Libraries were sequenced using 75bp single read output run on illumina NEXTseq platform.

Reads were trimmed using cutadapt[3] and mapped to mm10 mouse genome using STAR[4] v2.4.2a (default parameters). The pipeline quantifies the genes annotated in RefSeq (that have expanded with 1000 bases toward 5' edge and 100 bases toward 3' bases): /home/labs/bioservices/services/ngs/support/gtf/mm10.genes.3utr.gtf. Counting was done using htseq-count[5] (union mode). Further analysis is done for genes having a minimum of 5 read in at least one sample. Normalization of the counts and differential expression analysis was performed using DESeq2[6] with the following parameters: betaPrior=True, cooksCutoff=FALSE, independentFiltering=FALSE. Raw P values were adjusted for multiple hypothesis testing using the procedure of Benjamini and Hochberg. Pipeline was constructed using Snakemake[7].

mRNA-seq for the second biological repeat was done in parallel to ribosome footprint sequencing by a tailored mRNA-sequencing protocol as previously described[8]. Cells were harvested with Tri-Reagent (Sigma-Aldrich), total RNA was extracted, and poly-A selection was performed using Dynabeads mRNA DIRECT Purification Kit (Invitrogen). mRNA samples were subjected to DNAseI treatment and 3' dephpsphorylation using FastAP Thermosensitive Alkaline Phosphatase (Thermo Scientific) and T4 PNK (NEB) followed by 3' adaptor ligation using T4 ligase (NEB). The ligated products used for reverse transcription with SSIII (Invitrogen) for first strand cDNA synthesis. The cDNA products were 3' ligated with a second adaptor using T4 ligase and amplified for 8 cycles in a PCR for final library products of 200-300bp. Sequencing reads were aligned as previously described[9].

#### Ribosome footprint sequencing

For Ribo-seq libraries, cells were treated with 100µg/mL cycloheximide (CHX) for 1 minute. Cells were then placed on ice, washed twice with PBS containing 100µg/mL CHX, and lysed with lysis buffer (1% triton in 20mM Tris 7.5, 150mM NaCl, 5mM MgCl2, 1mM dithiothreitol supplemented with 10 U/ml Turbo DNase and 100µg/ml cycloheximide). After lysis, samples were flash-frozen and stored in -80°C until further processing. Subsequent, Ribo-seq library generation was performed as previously described[8]. Sequencing reads were aligned as previously described[9]. For comparing transcript expression level, mRNA and footprint counts were normalized to units of RPKM in order to normalize for gene length and for sequencing depth, based on the total number reads. Translation efficiency was calculated as the ratio between read counts (RPKM) of ribo-seq protected RNA fragments and mRNA sequencing.

#### tRNA sequencing and read analysis

tRNA sequencing protocol was adapted from Zheng et al., 2015 (Zheng et al., 2015) with minor modification. The protocol involves the use of a highly persistent and thermo-stable reverse transcriptase. The primers are DNA-RNA hybrids that have overhang of thymidine, which enable reverse transcription of adenosine ending RNAs. This leads to enrichment of mature tRNAs, due to their shared CCA tail. We isolated small RNAs of (<200bp) from the total RNA using SPRI-beads (Agencourt AMPure XP, Beckman Coulter; A63881), using dual side size-selection protocol. First RNA and beads were mixed at 1:1.8 ratio and supernatant was collected. The small RNA was isolated by mixing the clear supernatant with bead at 1:0.8 ratio, with addition of X1.34 isopropanol. Reverse transcription was done using TGIRT™-III Enzyme (InGex, LLC), with the indicated primers. 3' adaptor was ligated to the cDNA using T4 ligase (NEB; M0202S). The cDNA was purified using Dynabeads myOne SILANE (life Technologies; 37002D) after each step. The library was amplified using NEBNext PCR mix and cleaned using SPRI-beads. Samples were pooled and sequenced using a 75bp single read output run on MiniSeq high output reagent kit.

| **Primer name** | **Primer sequence** |
|---|---|
| Reverse-transcription DNA primer | 5'- CACGACGCTCTTCCGATCTT -3' |
| Reverse-transcription RNA primer | 5'-rArGrArUrCrGrGrArArGrArGrCrGrUrCrGrUrG -3' |
| 3'-ligation adaptor | 5'- AGATCGGAAGAGCACA-3' |

Read were trimmed using homerTool[10]. Alignment to the genome and mature tRNAs gene sequence was done using Bowtie2[11] with parameters --very-sensitive-local. Reads aligned with equal alignment score to the genome and mature tRNAs were annotated as mature tRNAs. Reads aligned to multiple tRNA genes were randomly assigned when mapping to identical anticodons, and discarded from the analysis if aligned to different anticodons. Read count was done using BedTools-coverage count[12]. Variant calling for detection of mutation and indels was done using samtools[13] command "mpileup" with the parameters: " -A -q1 -d100000".

Using this tRNA sequencing protocol, 45%-60% (depending on the sample) out of the reads are mapped to mature tRNAs, the cytosolic or mitochondrial. Additional 1%-6% of the reads were aligned to pre-mature tRNAs. Out of 273 unique mature tRNA sequences in the mouse genome, 120-156 tRNA genes get sufficiently high read counts that allow their quantification. On the other hand, 116-141 tRNAs did not get sufficient number of reads (under 20 reads). All the tRNA genes with a low scores ( below 50, tRNA-Scan SE[14]), have low read counts, and can be considered as pseudo-tRNA (Fig. S5-1A). The low read count indicates that those RNAs are either not expressed or cannot pass through the maturation process. Individual tRNAs were next grouped by anticodon. We calculated the gene copy number for each anticodon based on the number of tRNA genes in the genome, with prediction score above 50. We found that although they exist in the genome, tRNAs of certain anti-codons, with low gene copy number, have very low expression (Fig. S5-1B). For all of those anticodons, the matching codons can be translated by wobble interactions with a different tRNA group. These results are similar to measurements done in human HEK293 cells and in the brain[15], and they are in agreement with the predication that approximately a third of the tRNA genes in mouse are inactive in all tissues, and around 30% are tissue specific[16].

#### qRT-PCR for tRNA

small RNAs of (<200bp) were isolated from the total RNA using SPRI-beads (Agencourt AMPure XP, Beckman Coulter; A63881), by dual side size-selection protocol. Reverse transcription was done using TGIRT™-III Enzyme (InGex, LLC), with the indicated primers as in tRNA-sequencing protocol. Expression levels of $tRNA^{Val}$(AAC), $tRNA^{Gly}$(GCC) and $tRNA^{iMet}$, was normalized to $tRNA^{His}$(GTG) (constant expression based on tRNA sequencing). Levels were determined using quantitative RT-PCR with light cycler 480 SYBR I master kit (Roche Applied Science) and the LightCycler 480 system (Roche Applied Science), according to the manufacturer's instructions.

The primer used were:

| Primer name | Primer sequence |
|---|---|
| Mus-HisGTG2 Forward | GCCGTGATCGTATAGTGGTTAG |
| Mus-HisGTG2 Reverse | GTGCCGTGACTCGGATT |
| Mus-GlyGCC2 Forward | CATTGGTGGTTCAGTGGTAGAA |
| Mus-GlyGCC2 Reverse | CATTGGCCGGGAATCGAA |
| Mus-ValAAC2 Forward | GTTTCCGTAGTGTAGTGGTCATC |
| Mus-ValAAC2 Reverse | TGTTTCCGCCCGGTTTC |
| Mus-iMet Forward | CGCAGCGGAAGCGTG |
| Mus-iMet Reverse | GCAGAGGATGGTTTCGATCC |

#### LC/MS shotgun analyses of tRNAs

Capillary liquid chromatography/mass spectrometry (LC/MS) shotgun analyses of tRNA fraction were conducted as [17,18]. Total RNAs extracted from the culture cells were resolved by 10% PAGE with 7M urea. The gel bands corresponding to class I tRNA fraction were excised, followed by elution, the tRNA fraction was precipitated by ethanol. Two pmol of class I tRNA fraction was digested with 50 units RNase T1 (Thermo Scientific) in 20 mM $NH_4OAc$ (pH 5.3) at 37°C for 1h, followed by addition of an equal volume of 0.1 M triethylamine acetate (TEAA; pH 7.0). The digests were subjected to the L-column2 (C18, 5µm,ID 0.3× 5 mm, CERI) for desalting and chromatographed by HiQ sil C18W-3 nanospray column (C18, 3µm, 120 Å pore size, ID 0.1 × 100 mm, Techno Alpha) with solvent system consisted of 0.4 M 1,1,1,3,3,3-hexafluoro-2-propanol (HFIP) (pH 7.0) (solvent A) and 0.4 M HFIP (pH 7.0) in 50% methanol (solvent B) at a flow rate of 300 nl per min with a linear gradient of 5–100% B solvent over 35 min with a splitless nano HPLC system (DiNa, Techno Alpha). The eluent was ionized by ESI source and introduced into an ion trap-orbitrap hybrid mass spectrometer (LTQ Orbitrap XL, Thermo Fisher Scientific). Deprotonated ions of RNA fragments were detected with a negative polarity mode and scanned over an *m/z* range of 600–2000 throughout the separation. Extracted ion chromatograms (XICs) were plotted according to the theoretical *m/z* of each fragment.

#### Supplementary Figures

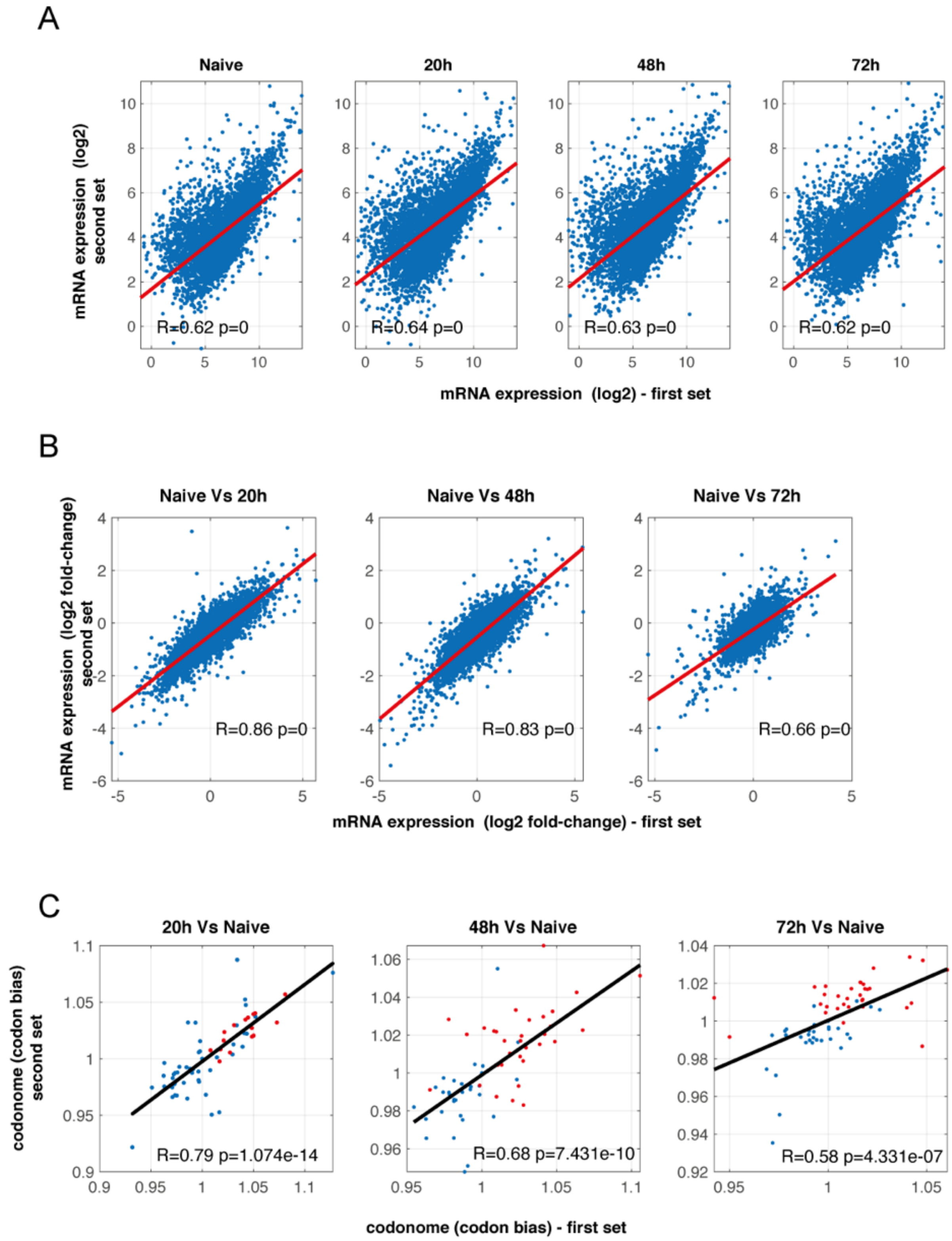


**Sup Figure 1-1- Comparisons of mRNA sequencing results of two biological independent experiments.** The first set is composed of six biological conditions (naïve, 20h, 48h CD62L$^+$, 48h CD62L$^-$, 72h CD62L$^+$, 72h CD62L$^-$) with 3 biological repeats for each condition, and the second from 4 biological conditions (naïve, 20h, 48h, 72h) with 2 biological repeats for each condition. For comparison between the sets CD62L$^+$ and CD62L$^-$ in the first set were averaged. A. Comparison of read counts (RPKM) for each mRNA. B. mRNA expression fold-changes (log2) between the indicated samples. C. Changes in codon usage, calculated based on mRNA expression (see M&M). marked in blue are G/C ending codons, marked in red are A/T ending codons.

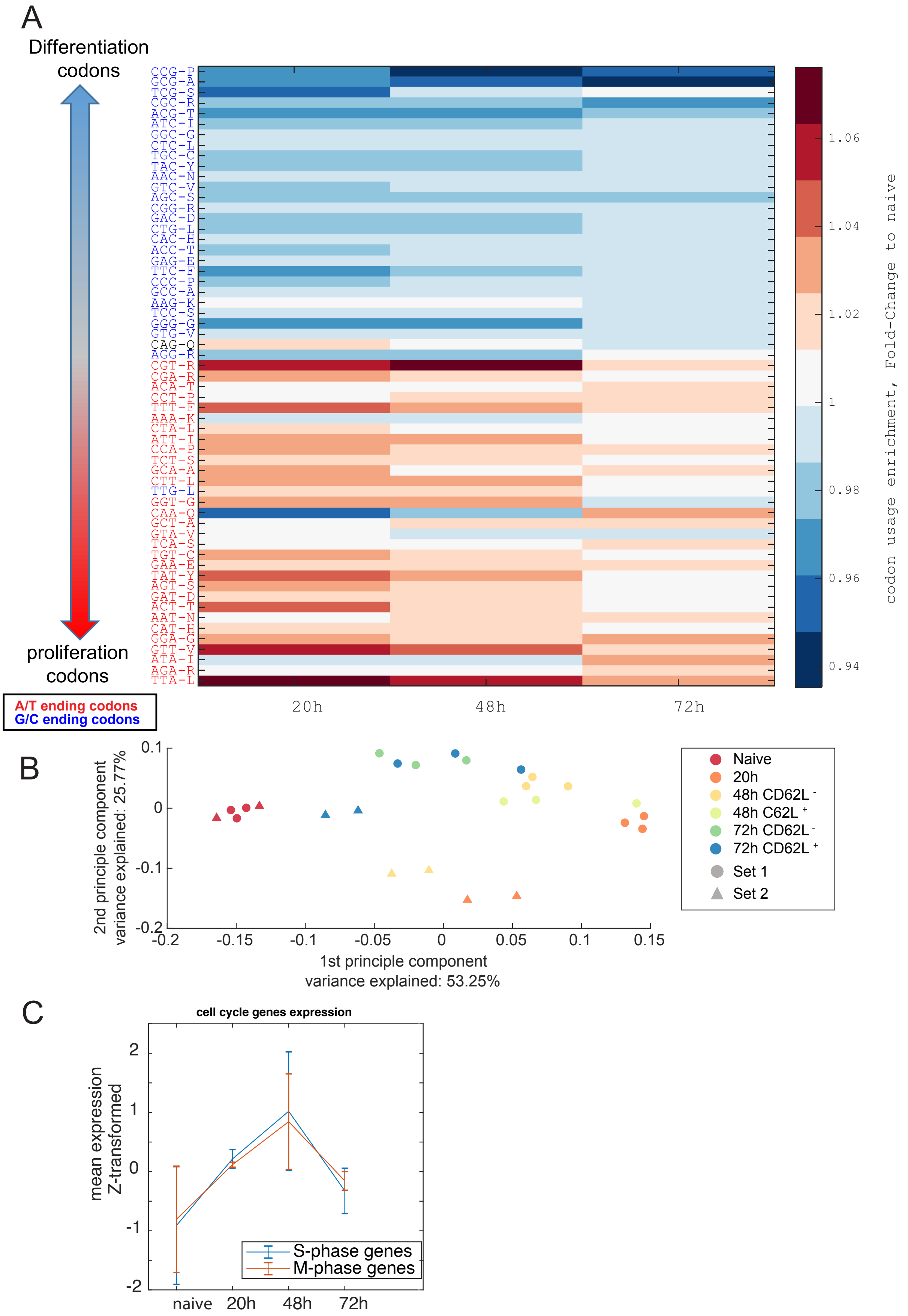
A
Differentiation codons
proliferation codons
CCG-P
GCG-A
TCG-S
CGC-R
ACG-T
ATC-I
GGC-G
CTC-L
TGC-C
TAC-Y
AAC-N
GTC-V
AGC-S
CGG-R
GAC-D
CTG-L
CAC-H
ACC-T
GAG-E
TTC-F
CCC-P
GCC-A
AAG-K
TCC-S
GGG-G
GTG-V
CAG-Q
AGG-R
CGT-R
CGA-R
ACA-T
CCT-P
TTT-F
AAA-K
CTA-L
ATT-I
CCA-P
TCT-S
GCA-A
CTT-L
TTG-L
GGT-G
CAA-Q
GCT-A
GTA-V
TCA-S
TGT-C
GAA-E
TAT-Y
AGT-S
GAT-D
ACT-T
AAT-N
CAT-H
GGA-G
GTT-V
ATA-I
AGA-R
TTA-L
20h
48h
72h
1.06
1.04
1.02
1
0.98
0.96
0.94
codon usage enrichment, Fold-Change to naive
A/T ending codons
G/C ending codons
B
2nd principle component variance explained: 25.77%
1st principle component variance explained: 53.25%
0.1
0
-0.1
-0.2
-0.2
-0.15
-0.1
-0.05
0
0.05
0.1
0.15
Naive
20h
48h CD62L -
48h C62L +
72h CD62L -
72h CD62L +
Set 1
Set 2
C
cell cycle genes expression
mean expression Z-transformed
2
1
0
-1
-2
naive
20h
48h
72h
S-phase genes
M-phase genes

**Sup Figure 1-2- changes in codon bias during T-cells activation. A.** Analysis of codon usage changes based on mRNA expression (normalized to amino acid, i.e. "codon-bias") as measured by RNA-seq. The analysis is similar to shown in Figure 1C, preformed on a biological independent experiment. Codons are sorted based on ratio of average codon usage of all genes from "G2/M phase" GO category to "pattern specification process" GO category. Separation between A/T (red) and G/C (blue) ending codons is significant in all comparisons ($p<10^{-5}$, rank-sum test). B. Codon bias differentiates proliferative and arrested T-cells. Shown here is a principal component analysis of the samples, based on their codon bias, as presented in panel A and Figure 1C (similar to Figure 1D yet, presented here are the two biological repeats). The first component (53% variance explained) best separates the naïve and 20h samples. Circles represent first biological repeat, triangles are the second biological repeat. C. Expression of S and M – phase genes in naïve and activated T-cells from second biological experiment (Z-transformed average of normalized read count of S-phase pathway (147 genes) and M phase pathway (326 genes), n=2, mean ± std).

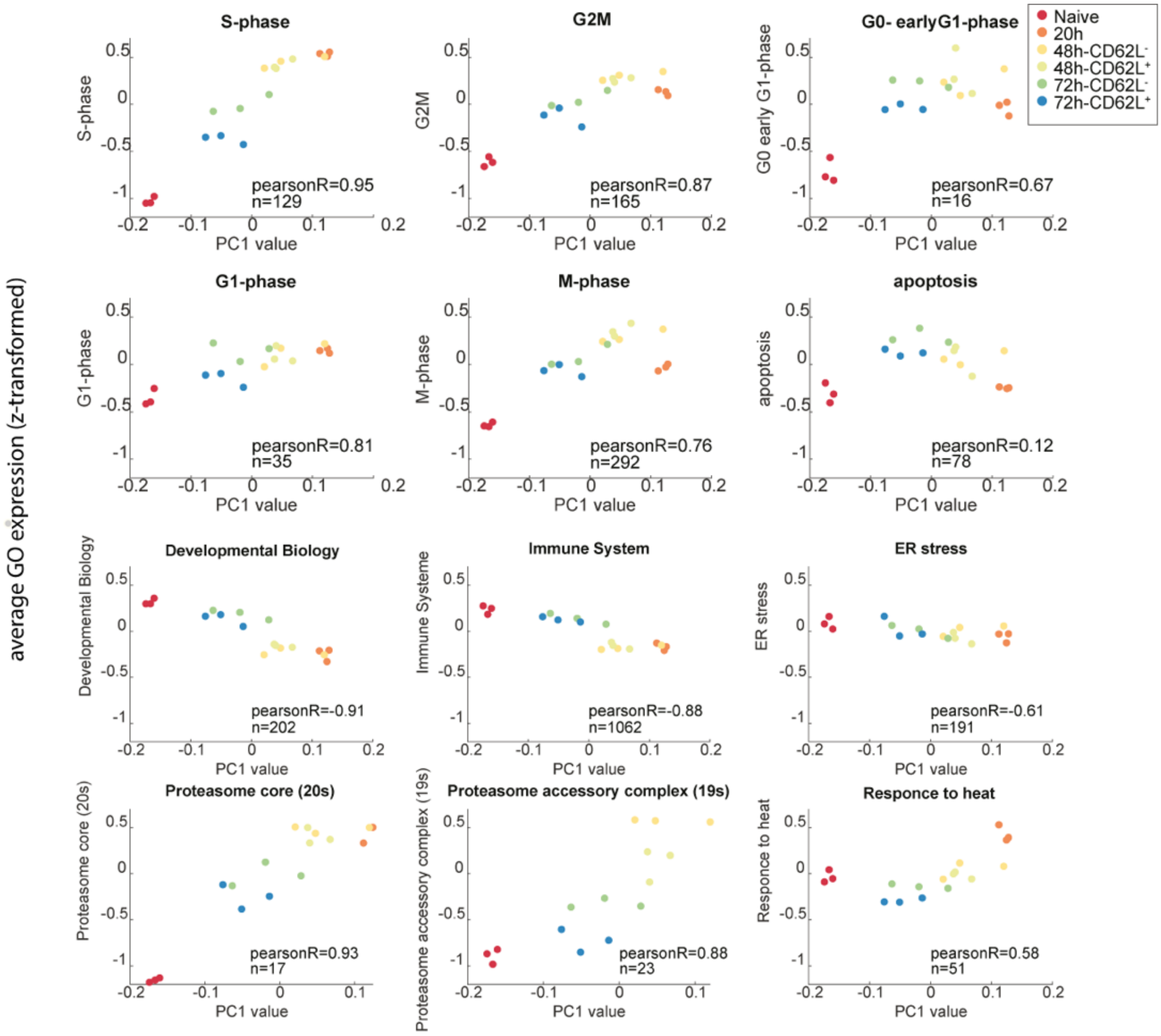

**Sup Figure 1-3 – Expression of GO related genes.** Scatter plots of position in PC1 (x-axis, calculated based on expressed codon usage, as in Figure 1C) plotted with average expression of the genes in the indicated GO category (y-axis, Z-transformed average of n genes) Each dot represent a sample. Each plot represent the indicated GO category.

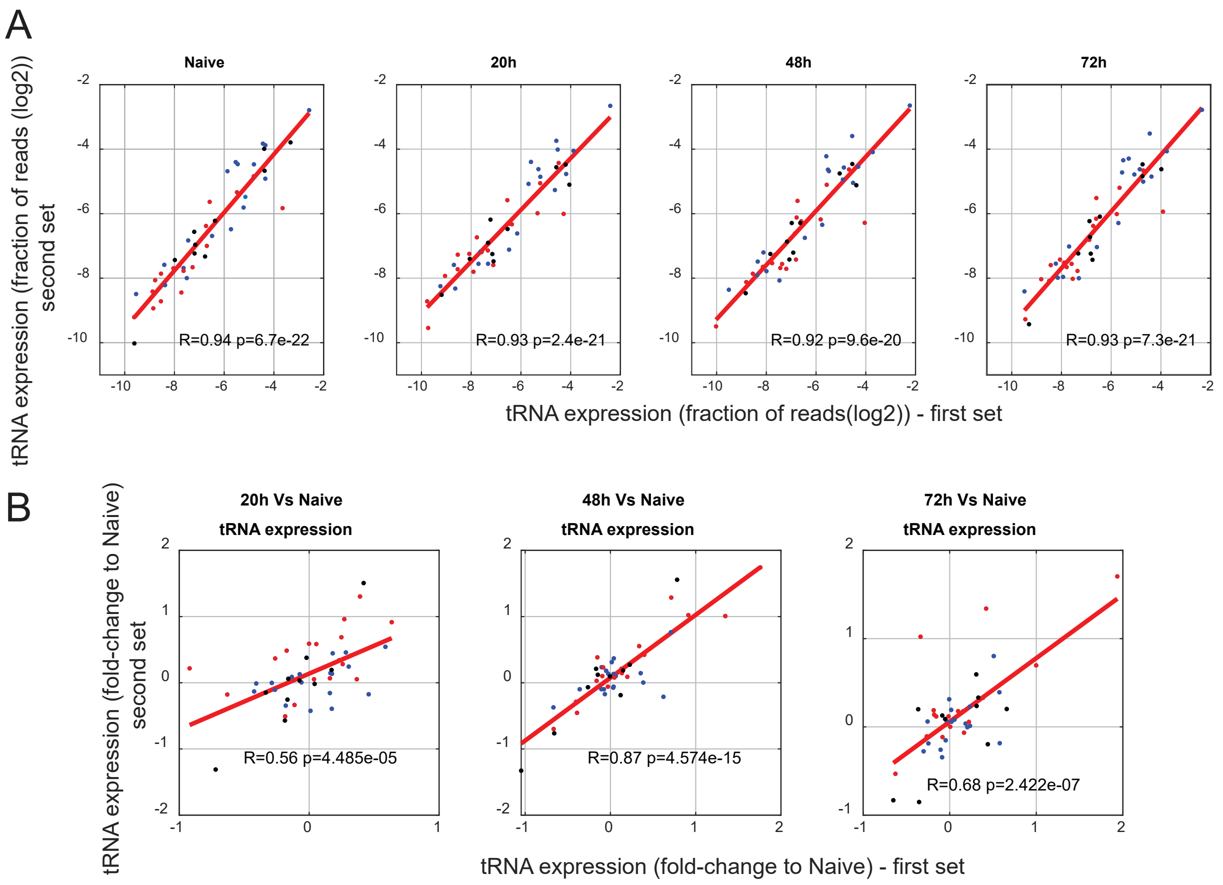


**Sup Figure 2-1- Comparisons of tRNA sequencing results of two biological independent experiments.** The first set is composed of six biological conditions (naïve, 20h, 48h CD62L$^+$, 48h CD62L$^-$, 72h CD62L$^+$, 72h CD62L$^-$) with 3 biological repeats for each condition, and the second from 4 biological conditions (naïve, 20h, 48h, 72h) with 3 biological repeats for each condition. For comparison between the sets CD62L$^+$ and CD62L$^-$ in the first set were averaged. A. Averaged tRNA expression summarized by anticodon (fraction of reads) of set 1 (x-axis) vs. set2 (y axis). (R and P for Pearson correlation, trend-line in red) B. tRNA expression fold-changes (log2) between the indicated samples matched in the two experimental sets (R and P for Pearson correlation, trend-line in red) red dots marks tRNA coding for A/T ending codons, Blue dots marks tRNA coding for G/C ending codons, black dots marks wobble/other tRNAs.

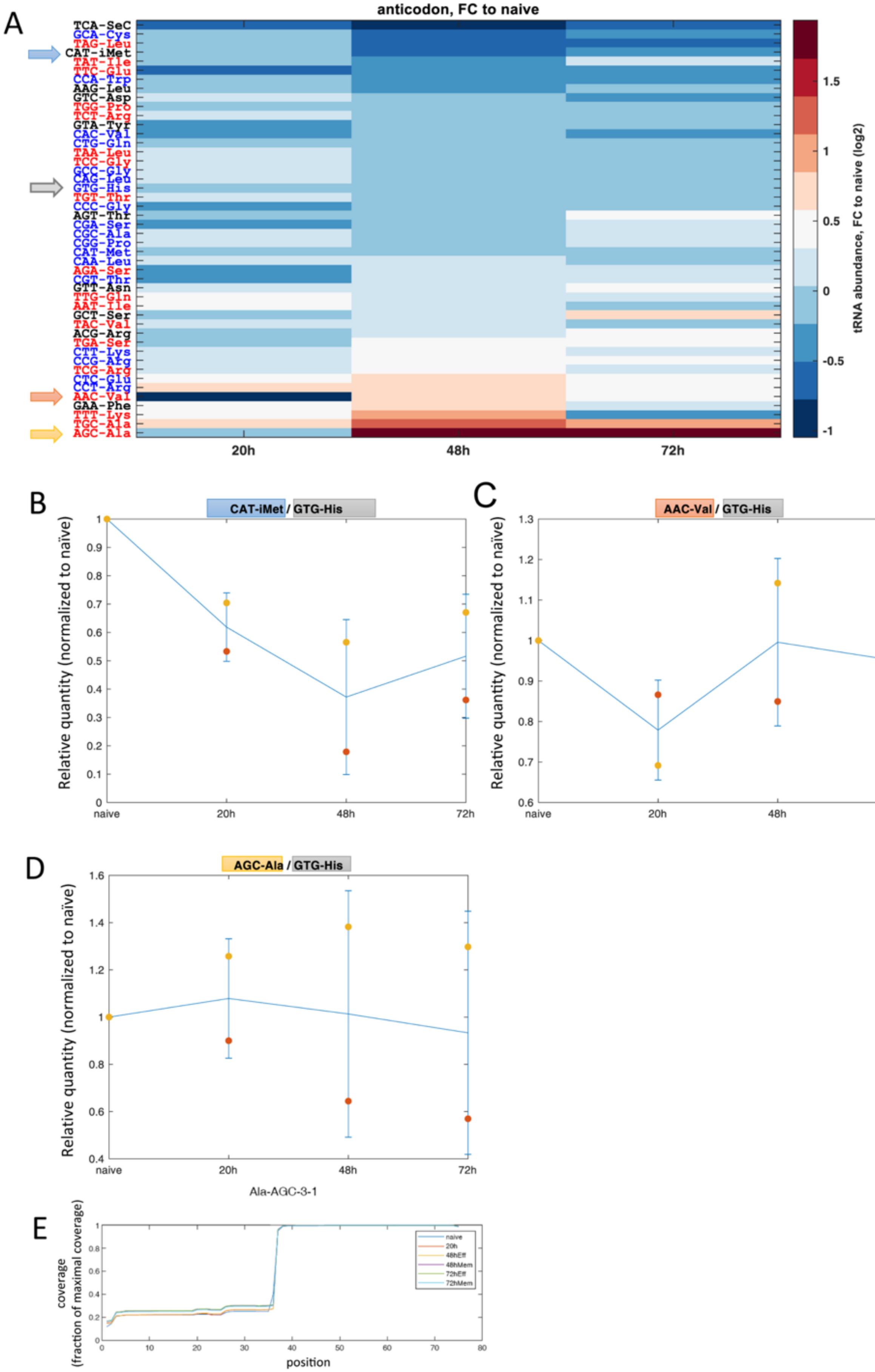
A
anticodon, FC to naive
TCA-SeC
GCA-Cys
TAG-Leu
CAT-iMet
TAT-Ile
TTC-Glu
CCA-Trp
AAG-Leu
GTC-Asp
TGG-Pro
TCT-Arg
GTA-Tyr
CAC-Val
CTG-Gln
TAA-Leu
TCC-Gly
GCC-Gly
CAG-Leu
GTG-His
TGT-Thr
CCC-Gly
AGT-Thr
CGA-Ser
CGC-Ala
CGG-Pro
CAT-Met
CAA-Leu
AGA-Ser
CGT-Thr
GTT-Asn
TTG-Gln
AAT-Ile
GCT-Ser
TAC-Val
ACG-Arg
TGA-Ser
CTT-Lys
CCG-Arg
TCG-Arg
CTC-Glu
CCT-Arg
AAC-Val
GAA-Phe
TTT-Lys
TGC-Ala
AGC-Ala
20h
48h
72h
1.5
1
0.5
0
-0.5
-1
tRNA abundance, FC to naive (log2)
B
CAT-iMet / GTG-His
Relative quantity (normalized to naïve)
naive
20h
48h
72h
C
AAC-Val / GTG-His
Relative quantity (normalized to naïve)
naive
20h
48h
72h
D
AGC-Ala / GTG-His
Relative quantity (normalized to naïve)
naive
20h
48h
72h
Ala-AGC-3-1
E
coverage
(fraction of maximal coverage)
position
naive
20h
48hEff
48hMem
72hEff
72hMem

**Sup Figure 2-2- Dynamics of tRNA expression during T-cell activation**. A. Heat-map representation of changes in tRNA expression at each time point, normalized by expression in Naïve T-cells. The analysis is similar to shown in Figure 2B, preformed on a biological independent experiment. Rows are sorted based on the tRNA expression fold-change between naïve and 48h after activation. Arrows indicate tRNA measured by qRT-PCR. B-D. qRT-PCR for the indicated tRNA, normalized by qRT-PCR of tRNA GTG-His. E. Coverage of reads aligned to tRNA-Ala-AGC at each position.

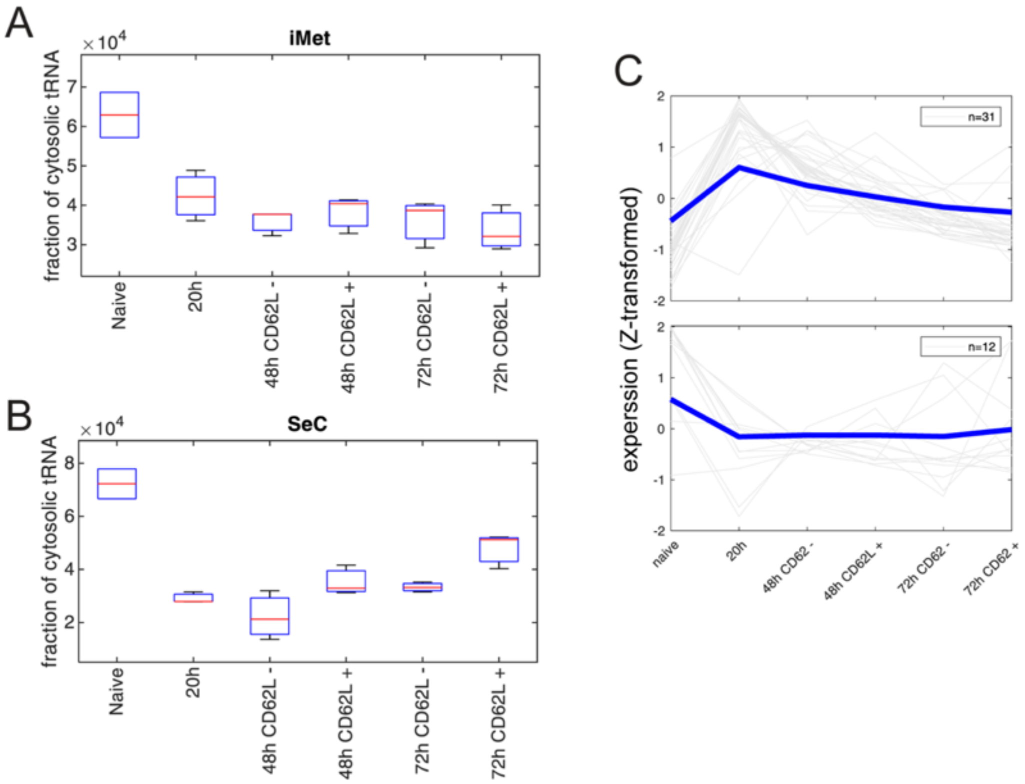


**Sup Figure 2-3- Expression of iMet and SeC tRNAs and translation initiation factors during T-cells activation process.** A. Expression of iMet-tRNA (fraction of cytosolic tRNA). B. expression of SeC tRNA (fraction of cytosolic tRNA). C. mRNA expression of translation initiation factors. Shown here mean of Z-transformed expression. Expression patterns are clustered into 2 main clusters- the lower panel (n=12 genes) shows genes with expression pattern correlative to iMet, while the upper panel has an opposing expression pattern (n=31).

A

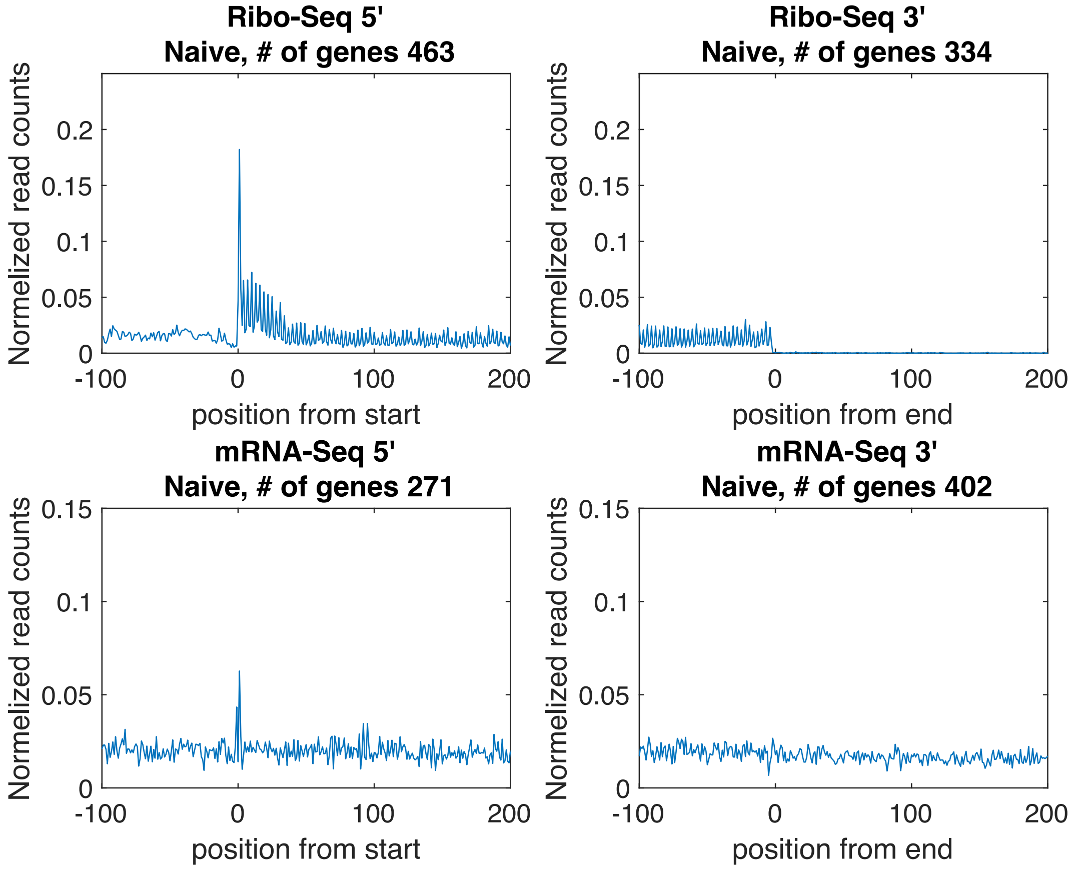


B

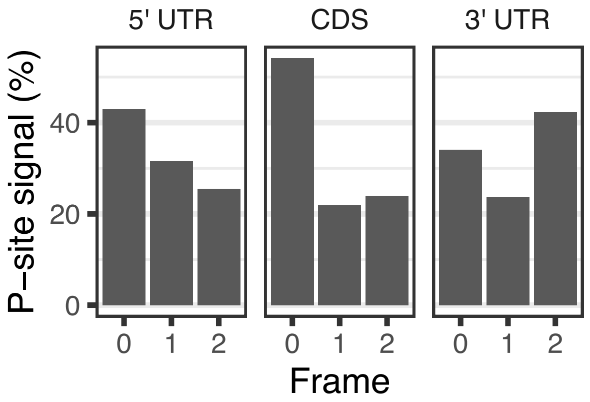


**Sup Figure 2-4- Ribo-seq quality assessments.** A. Plots showing reads density in ribo-seq (upper panel) and mRNA-seq (lower panel). The reads are align to the start codon (left) or the stop codon (right). The Ribo-seq reads are enriched at the start and stop codon, and are higher in the coding region compared to the UTR. B. Ribosome reading frame in 5' UTR, coding sequence and 3'UTR. Frame is calculated based on P site position assigned for each read according to ribosome protected read length using riboWalts[19] (see M&M).

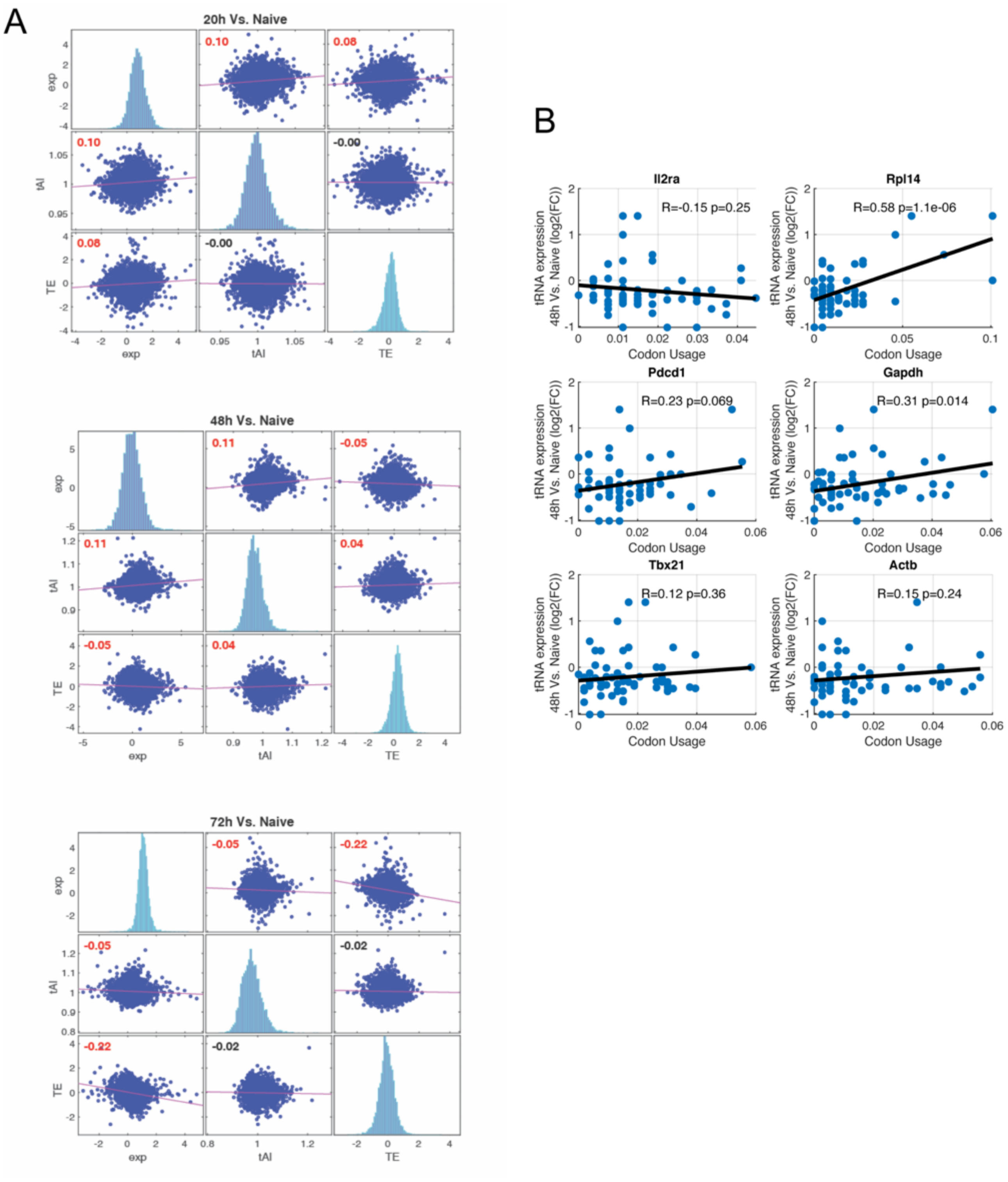


**Sup Figure 2-5- Gene level correlations- scatter plots of mRNA expression, tAI and TE.** Each dot represents a gene, the axes represent mRNA expression (log2FC), tAI (fold change to Naïve) and TE (log2FC). The different panels represent different time points during T-cell activation process relative to Naïve cells. Marked in red are significant correlations (p-value <0.05). B. correlations between codon usage of specific genes (left- selected T-cells related genes, right- highly expressed genes) with fold-change in tRNA expression of 48h vs. naïve T-cells (log2).

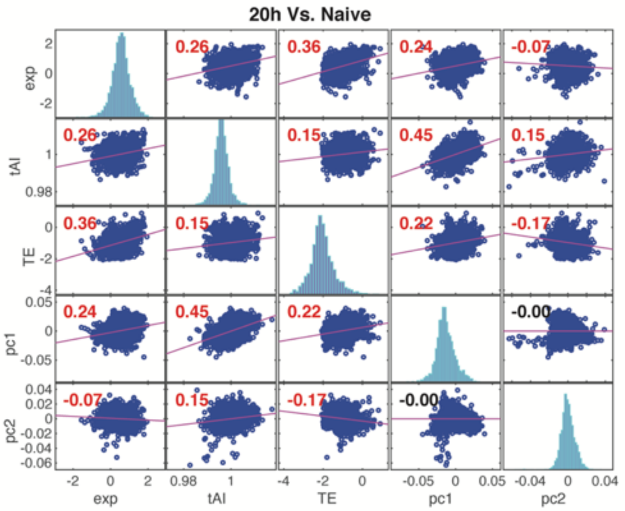
20h Vs. Naive
exp
tAI
TE
pc1
pc2

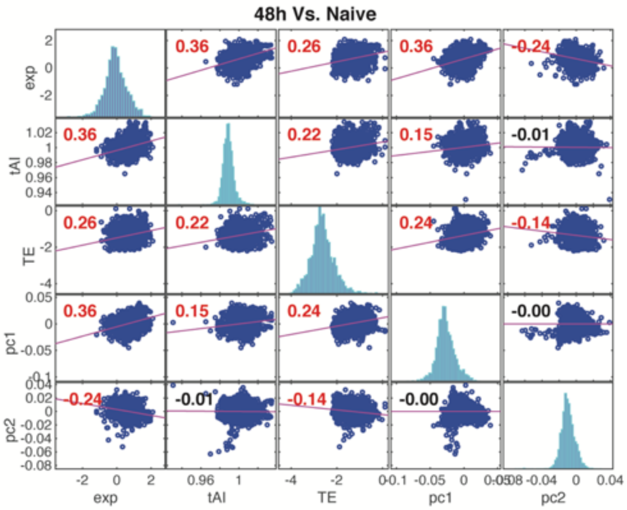
48h Vs. Naive
exp
tAI
TE
pc1
pc2

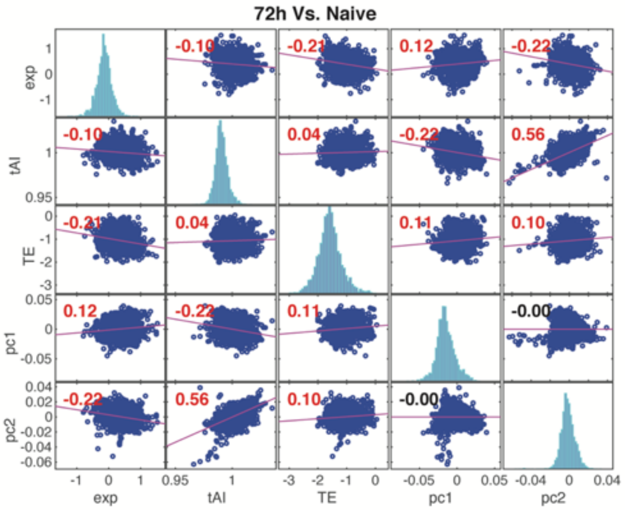
72h Vs. Naive
exp
tAI
TE
pc1
pc2

**Sup Figure 2-6- GO level correlations- scatter plots of expression, tAI, TE and location on the PCA presented in figure 2D.** Each dot represents a gene set derived from GO category, the axes represent the average value (Z-transformed) of expression, tAI and TE as well as correlation to 1st and 2nd PC calculated based on codon usage of GO categories (as in Figure 2D). The different panels represent different time points during T-cell activation process relative to Naïve cells. Marked in red are significant correlations (p-value <0.05).

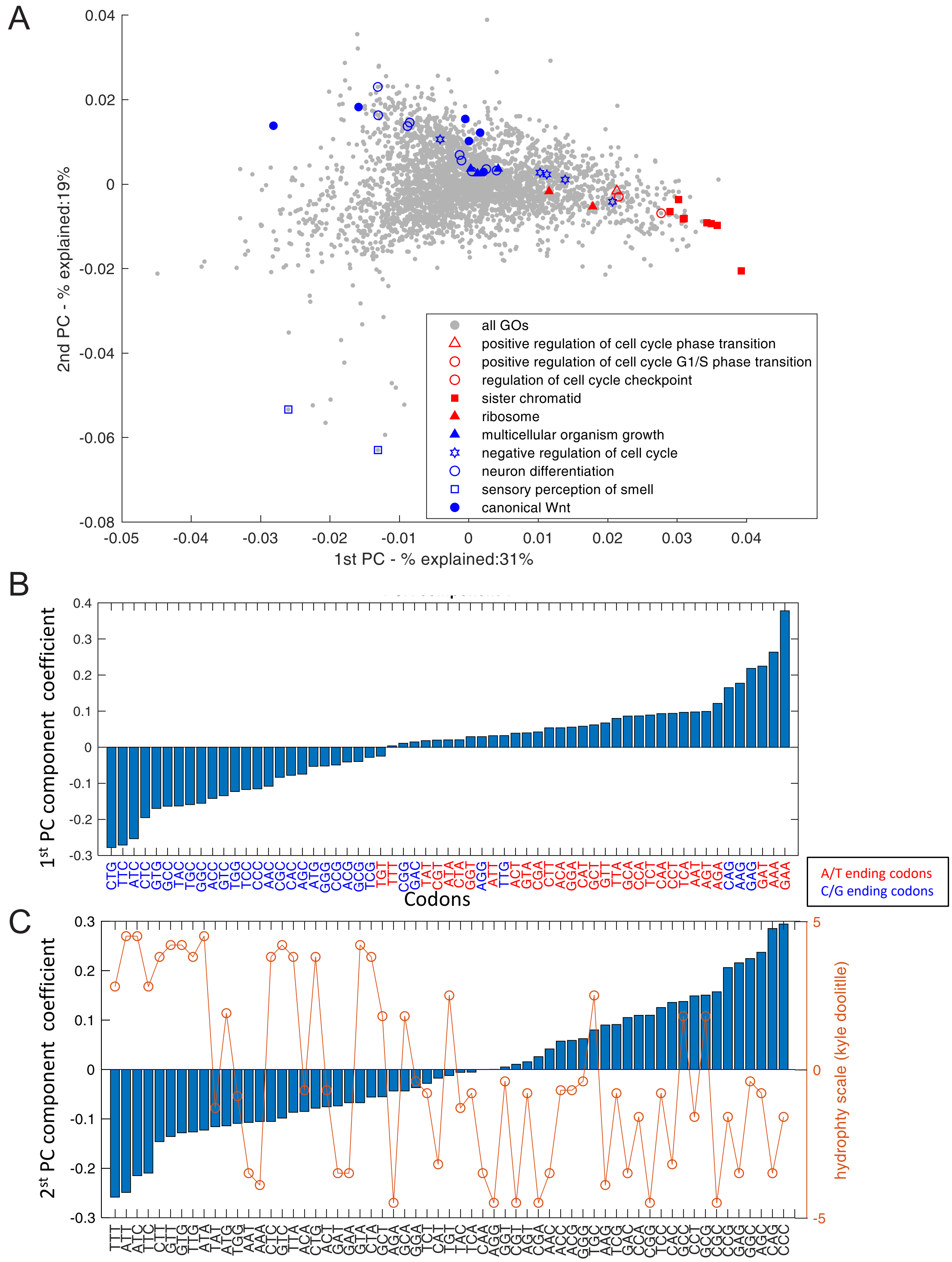

**Sup Figure 2-7- A. PCA projection of the mouse codon usage of gene sets derived from GO categories.** A. The location of each gene set in this space is determined by the average codon usage of all the genes that belong to it. The % variance, out of the total original variance in the high-dimensional space, spanned by the first and second PCs is indicated on the x and y axis, respectively. Marked in red are gene categories related to cell autonomous functionality (proliferation related), marked in blue are GO categories related to multi-cellularity functions (tissue specific) B. codon coefficient contribute to the 1st PC. Marked in red codons end with A/T, marked in blue codons that end with G/C. C. codon coefficient contribute to the 2nd PC. In orange is the kyte-doolittle hydropathy score of the corresponding amino-acid[20]

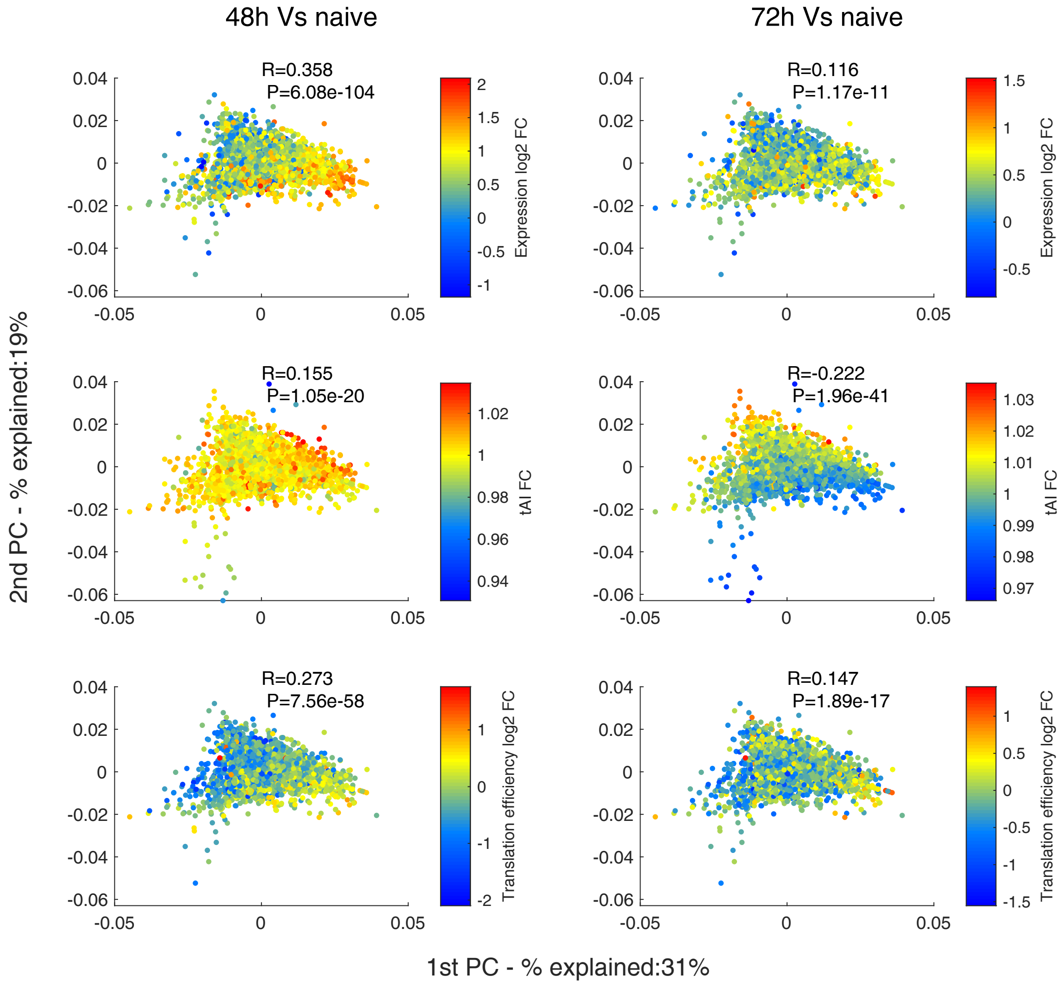


**Sup Figure 2-8-** A PCA projection of the mouse codon usage of gene sets derived from GO categories. Each point represents one gene set. Gene sets corresponding to tissue-specific GO terms are to the left side, and those corresponding to proliferation related GO terms are to the right. The color code in the upper panel represent changes at the mRNA level, averaged over all the genes in each gene set. In the middle panel, each gene category is color coded according to the relative change in availability of the tRNAs that correspond to the codon usage of its constituent genes, averaged over all genes in the gene set. A red color for a given gene set indicates that on average the genes in that set have codons that mainly correspond to the tRNAs that are induced in the condition, whereas a blue color indicates that the codon usage in the set is biased toward the tRNAs that were repressed in that given condition. In the lower panel the color code indicates translation efficiency (TE) calculated based on ribo-seq by ratio of reads of the ribosome protected fragment (RPKM) divided by read count of mRNA-seq (RPKM). Right panels show 48h vs. Naïve, left panels show 72h vs. naïve.

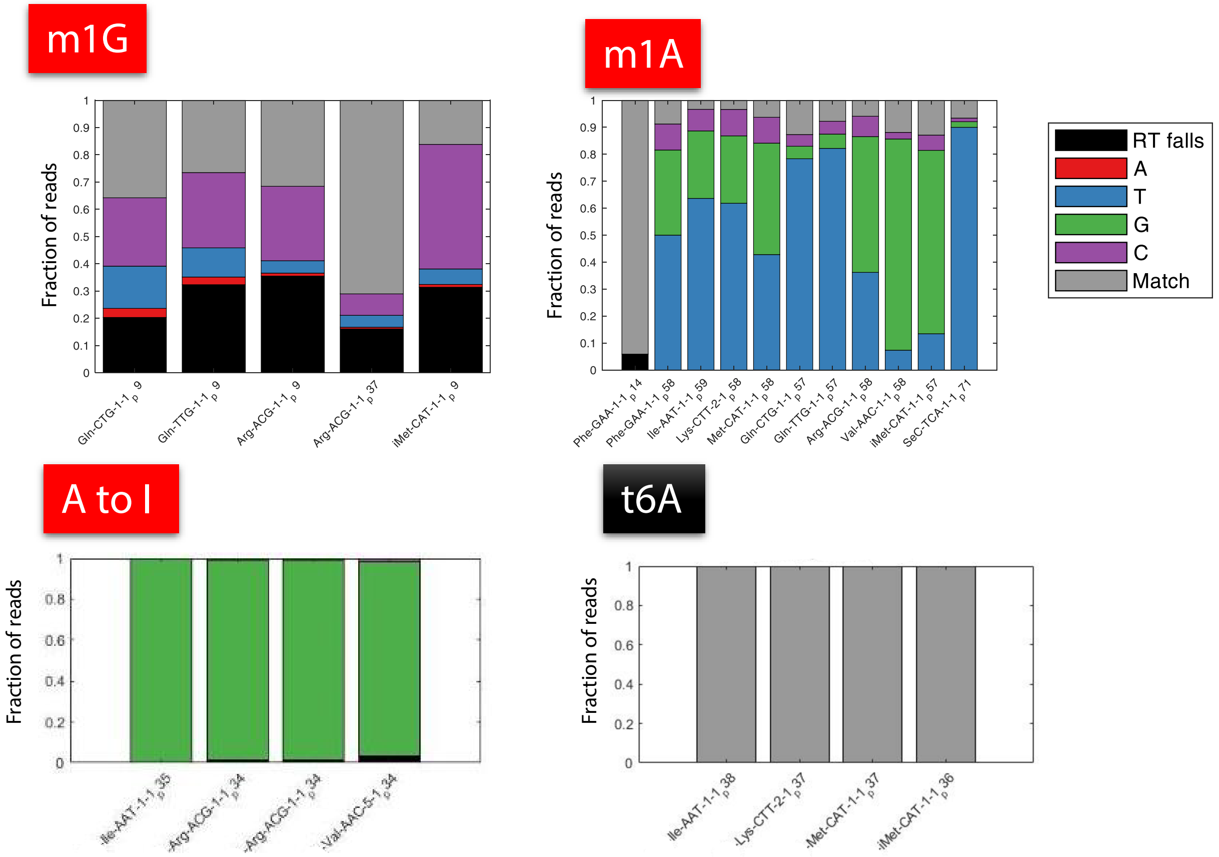


**Sup Figure 3-1- modification patterns identified by tRNA sequencing**. Each plot represent the indicated modification type, each bar is an annotated modification position based on MODOMICS database for mouse tRNA. Colors indicated mismatch to the indicated base in the alignment, black indicate RT-abortion, and gray indicated a matched read.

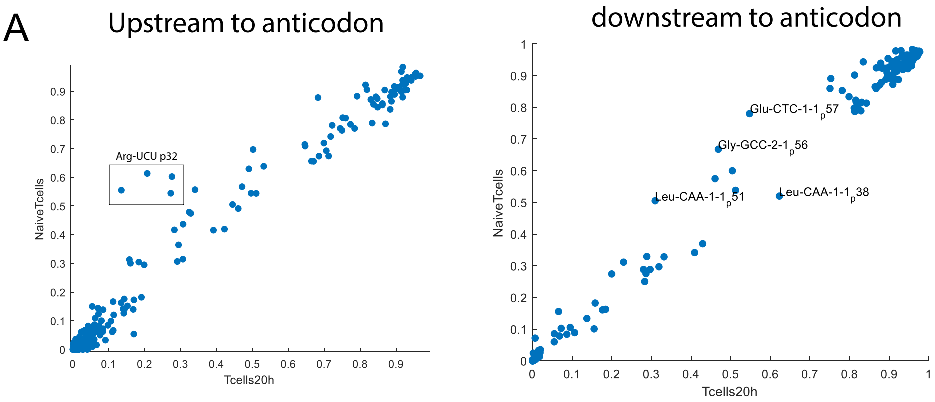


**Sup figure 3-2- modifications detection based on tRNA sequencing.** A. average modification levels base on tRNA-seq from mice naïve and activated (20h) T-cells (n=3). Threshold for modification calling is: minimum coverage of modified position = 20 reads, modification fraction above 10% in at least one sample. Left panel- upstream to anticodon, right panel- downstream to anticodon.

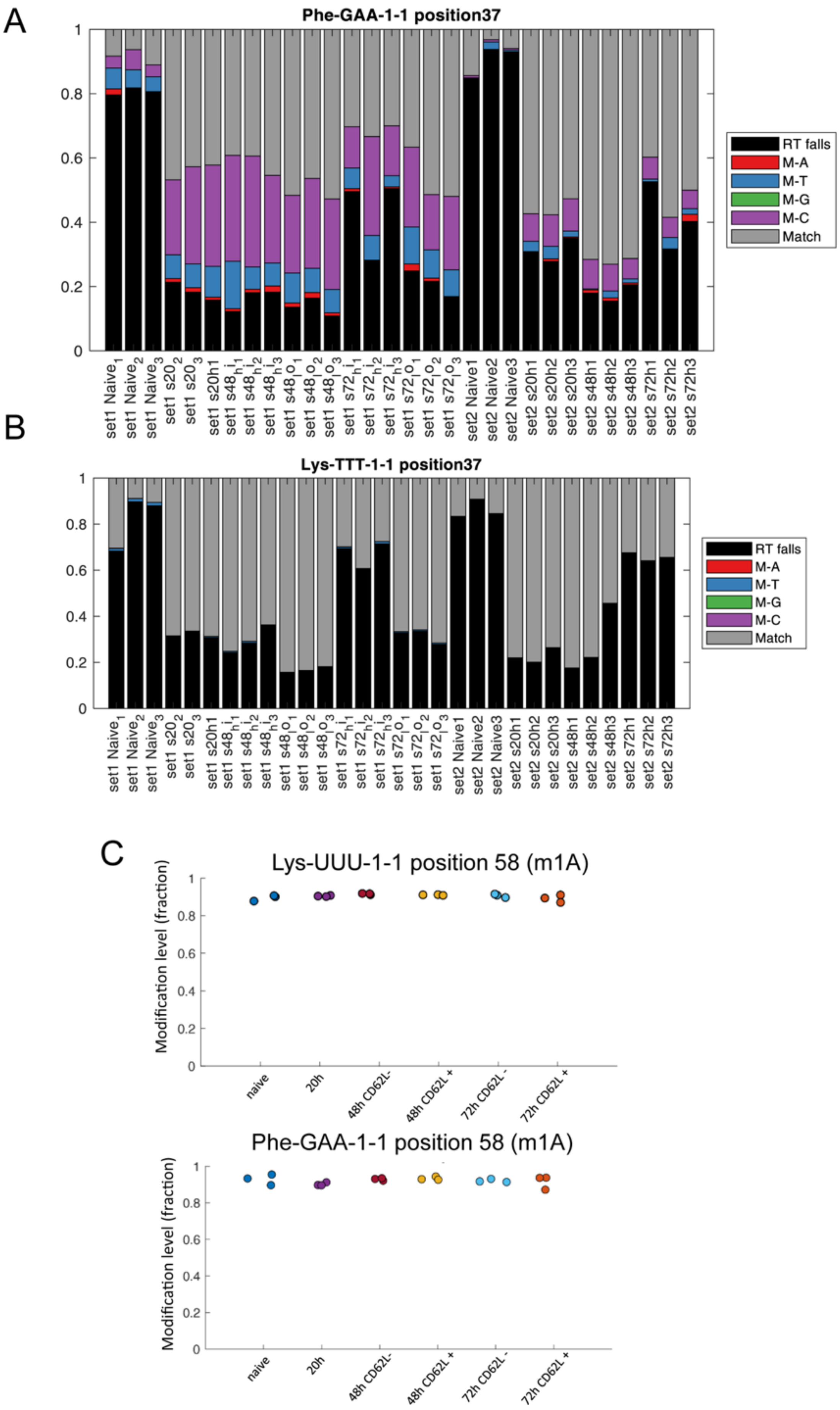
A
Phe-GAA-1-1 position37
RT falls
M-A
M-T
M-G
M-C
Match
set1 Naive1
set1 Naive2
set1 Naive3
set1 s20 2
set1 s20 3
set1 s20h1
set1 s48 h i 1
set1 s48 h i 2
set1 s48 h i 3
set1 s48 l o 1
set1 s48 l o 2
set1 s48 l o 3
set1 s72 h i 1
set1 s72 h i 2
set1 s72 h i 3
set1 s72 l o 1
set1 s72 l o 2
set1 s72 l o 3
set2 Naive1
set2 Naive2
set2 Naive3
set2 s20h1
set2 s20h2
set2 s20h3
set2 s48h1
set2 s48h2
set2 s48h3
set2 s72h1
set2 s72h2
set2 s72h3
B
Lys-TTT-1-1 position37
RT falls
M-A
M-T
M-G
M-C
Match
C
Lys-UUU-1-1 position 58 (m1A)
Modification level (fraction)
naive
20h
48h CD62L-
48h CD62L +
72h CD62L -
72h CD62L +
Phe-GAA-1-1 position 58 (m1A)
Modification level (fraction)

**Sup Figure 3-3- modification pattern detected by tRNA sequencing.** A. Bars show the mismatches and RT failures at position 37 in tRNA-Phe-GAA in all samples. Colors indicated mismatch to the in the alignment, black indicate RT-abortion, and gray indicated a matched read. B. Bars show the mismatches and RT failures at position 37 in tRNA-Lys-TTT in all samples. Colors indicated mismatch to the in the alignment, black indicate RT-abortion, and gray indicated a matched read. C. m1A modification at position 58 remains constant throughout the activation process in tRNA-Lys-UUU (upper panel) and tRNA-Phe-GAA (lower panel).

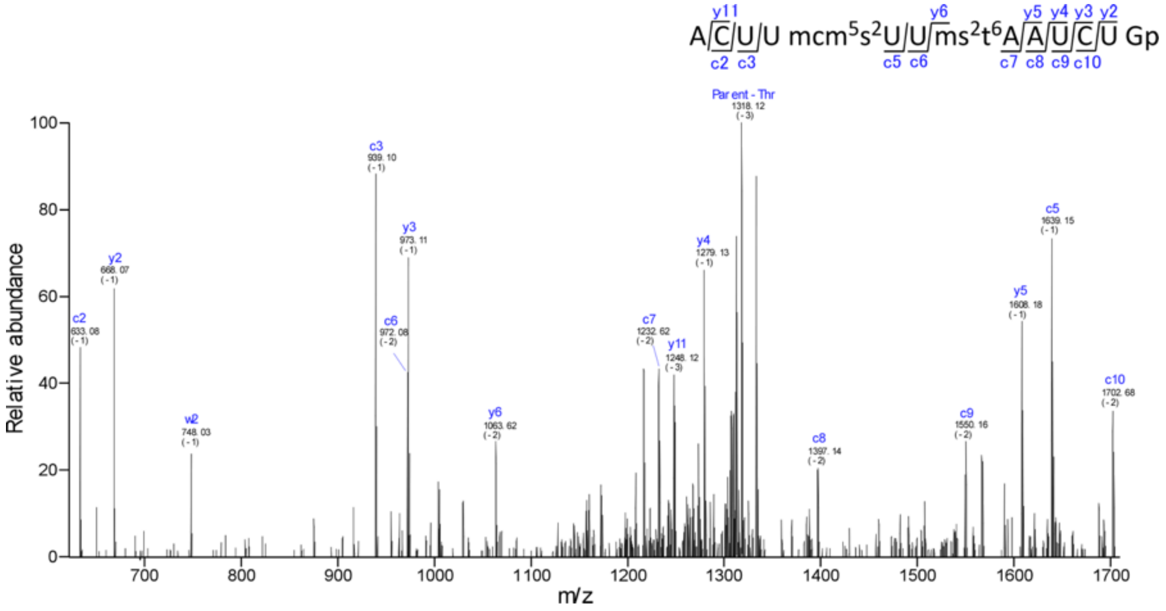


**Sup Figure 3-4- Collision-induced dissociation (CID) spectrum of the anticodon-containing fragment of cytoplasmic $tRNA^{Lys}$ in mouse T-cells.** The triply-charged negative ion of the anticodon-containing fragment (*m/z* 1358.15) bearing mcm5s2U34 and ms2t6A37 was used as a precursor ion for CID. The product ions were assigned as described [21]. Sequence of the parent fragment with assigned product ions are indicated in the inset.

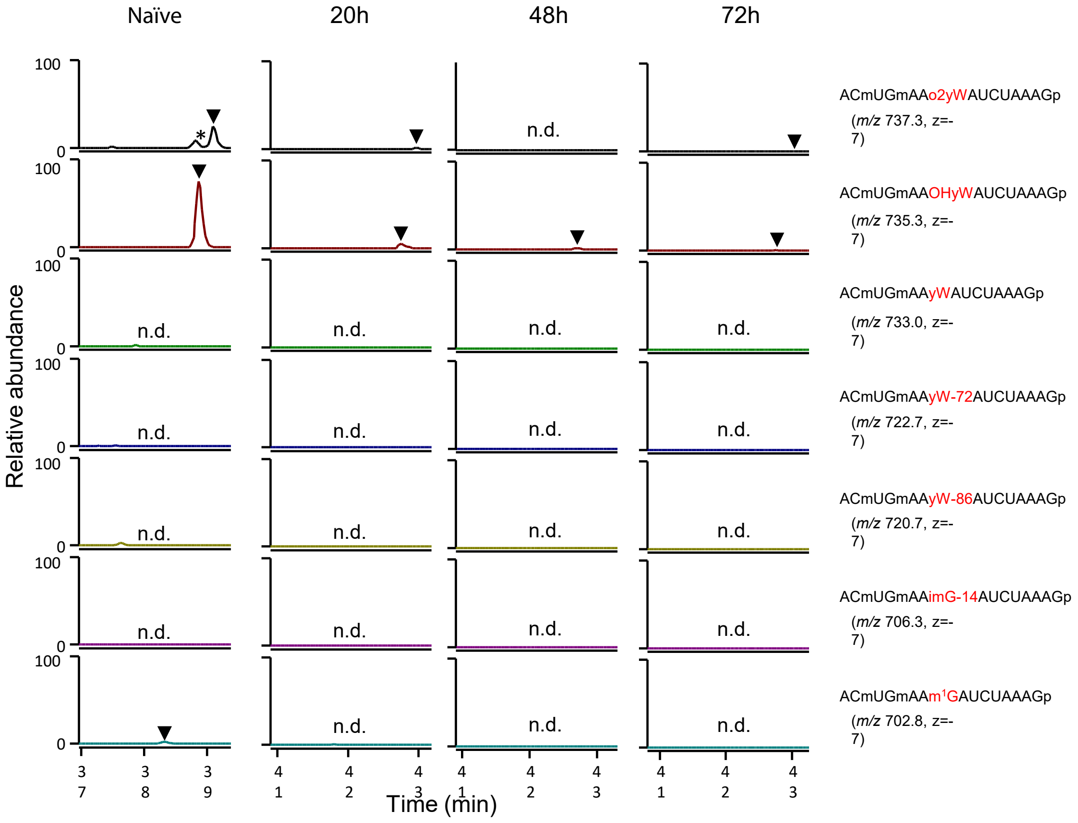


**Sup Figure 3-5- . Mass spectrometric shotgun analysis of cytoplasmic tRNA modifications during T-cell activation.** Shotgun analysis of class I tRNA fraction from mouse CD4+ T-cells collected at 0 (naive), 20, 48 and 72h after activation. The tRNA fraction was digested by RNase T1 and subjected to capillary LC/MS. Top to bottom panels represent XICs for negative ions of anticodon-containing fragments of cytoplasmic $tRNA^{Phe}$ bearing o2yW37, PHyW37, yW37, yW-72, yW-86, imG-14 and m1G, respectively. The peak heights in each condition are normalized by $AAAUAm^2Gp$ (*m/z* 665.4, z=-4) which is an internal fragment of cytoplasmic $tRNA^{Phe}$. n.d. stands for not detected. The actual peak is indicated by arrowhead. Non-specific peak is asterisked.

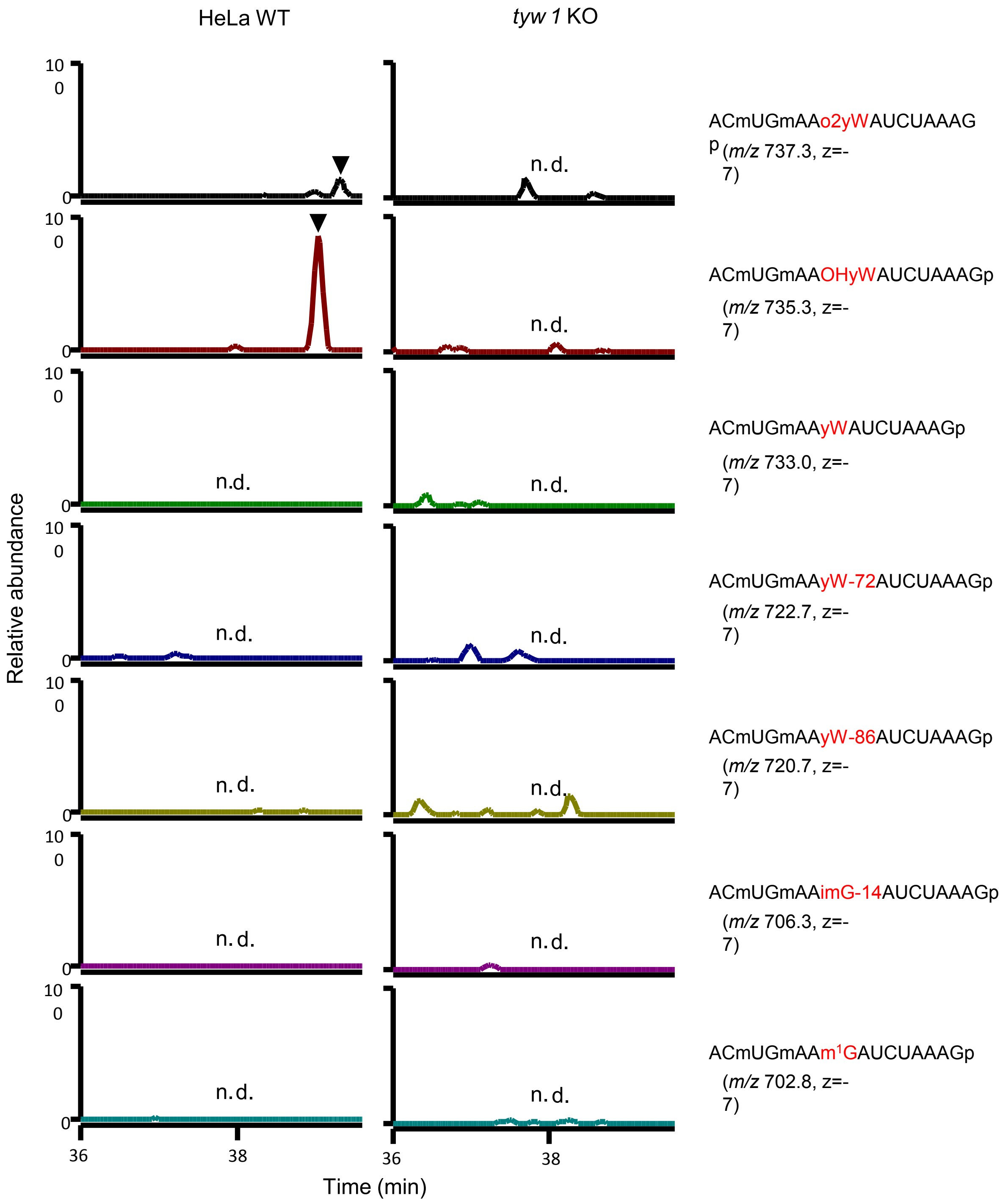

HeLa WT
tyw 1 KO
Relative abundance
100
0
n.d.
ACmUGmAAo2yWAUCUAAAGp (m/z 737.3, z=-7)
ACmUGmAAOHyWAUCUAAAGp (m/z 735.3, z=-7)
ACmUGmAAyWAUCUAAAGp (m/z 733.0, z=-7)
ACmUGmAAyW-72AUCUAAAGp (m/z 722.7, z=-7)
ACmUGmAAyW-86AUCUAAAGp (m/z 720.7, z=-7)
ACmUGmAAimG-14AUCUAAAGp (m/z 706.3, z=-7)
ACmUGmAAm1GAUCUAAAGp (m/z 702.8, z=-7)
36
38
Time (min)

**Sup Figure 4-1- Mass spectrometric shotgun analysis of cytoplasmic tRNA$^{Phe}$.** Shotgun analysis of class I tRNA fraction from HeLa WT (left panels) and *tyw1* KO (right panels) cells. The tRNA fraction was digested by RNase T1 and subjected to capillary LC/MS. Top to bottom panels represent XICs for negative ions of anticodon-containing fragments of cytoplasmic tRNA$^{Phe}$ bearing o2yW37, PHyW37, yW37, yW-72, yW-86, imG-14 and m1G, respectively. n.d. stands for not detected. The actual peaks are indicated by arrowheads.

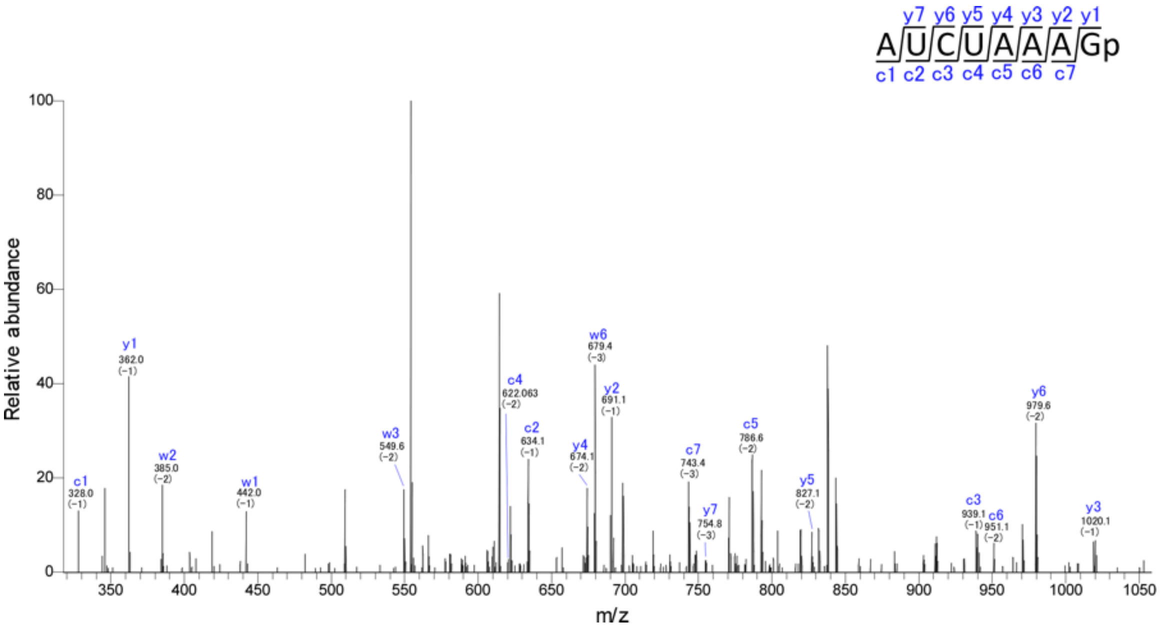


**Sup Figure 4-1- Collision-induced dissociation (CID) spectrum of the RNase $T_1$-digested fragment of cytoplasmic tRNA$^{Phe}$ in HeLa *tyw1* KO cells.** The quatruply-charged negative ion of AUCUAAAGp (*m/z* 648.1. z=-4) was selected as a precursor ion for CID. The product ions were assigned as described [21]. Sequence of the parent fragment with assigned product ions are indicated in the inset.

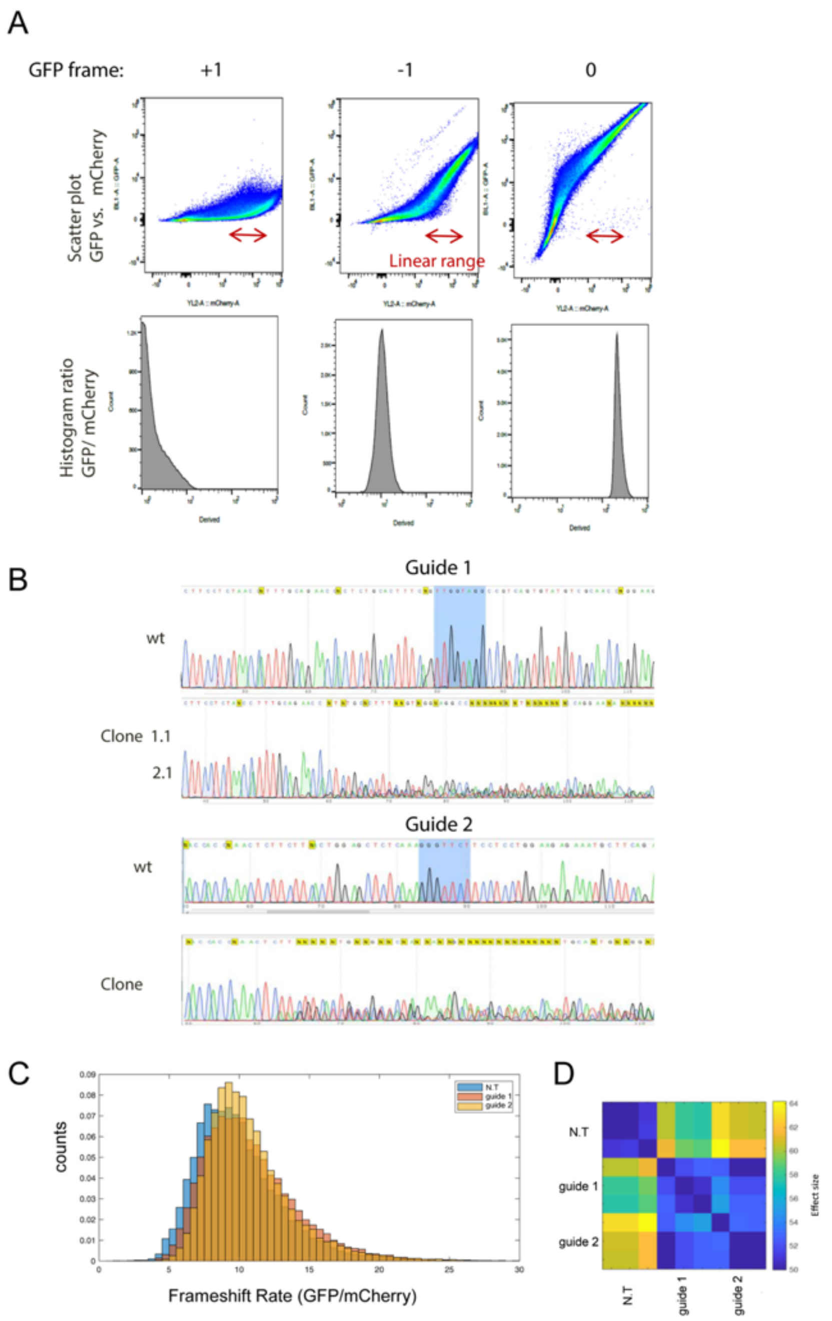
A
GFP frame:
+1
-1
0
Scatter plot GFP vs. mCherry
Linear range
Histogram ratio GFP/ mCherry
B
Guide 1
wt
Clone 1.1
2.1
Guide 2
wt
Clone
C
counts
Frameshift Rate (GFP/mCherry)
N.T
guide 1
guide 2
D
N.T
guide 1
guide 2
Effect size

**Sup Figure 4-3- Frameshift detection in *tyw1*-KO cells.** A. Upper panel are scatter plots of GFP (y-axis) and mCherry (x-axis) intensity measured in HeLa cells expressing mCherry- HIV frameshift signal – GFP reporter, in +1, -1 and 0 frame. Red arrows indicate the linear area we used to calculate the ratio between GFP and mCherry. Lower panel shows the ratio GFP/mCherry in the same cells as in upper panel. B. Validation of *tyw1* KO in HeLa cells. shown here are Sanger sequencing chromatogram of CRISPR edited cells using 2 different gRNA for *tyw1,* compared with non-targeting (N.T) gRNA*.* C. histograms of GFP/mCherry ratio (as in A.) measured in *tyw1* KO and N.T. HeLa cells. D. shown are the calculated effect size for all comparisons. All comparisons of tyw1 Vs. N.T were higher then 56, indicating a medium effect size (rank-sum p-value <$10^{-5}$).

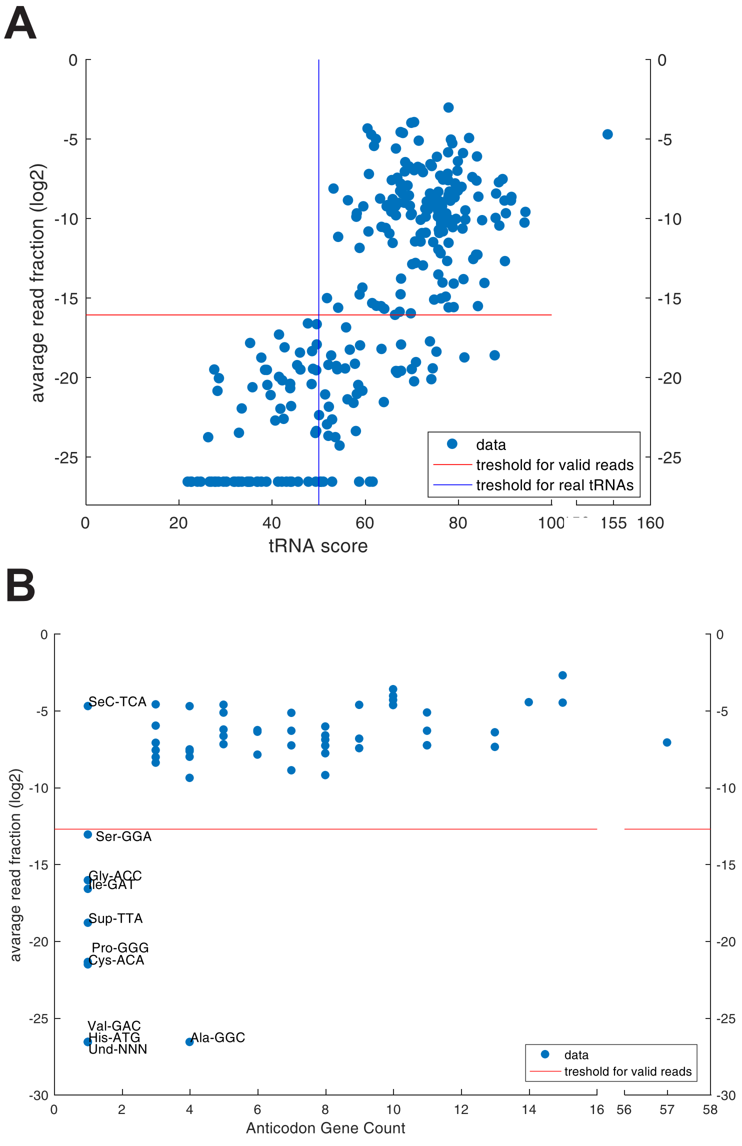


**Sup figure 5-1- tRNA Deep sequencing in T-cells shows agreement with codon usage and tRNA score.** A. tRNA read count is in agreement with tRNA score. Shown here is a scatter plot of the predicted tRNA score as calculated by tRNAscan-SE (ref) compare to fraction of reads of each unique tRNA. tRNAs with score below 50 consider to be pseudo-tRNAs, and indeed gets very few reads. B. anticodon read count. Shown here is a scatter plot of the tRNA gene copy number, calculated based on the number of tRNA of the same anti-codon with score above 50, compare to fraction of reads of each tRNA anti-codon group

#### Supplementary Tables

| Detectable modification | Undetectable modification |
| --- | --- |
| I | Ar(p) |
| N | Cm |
| acp3U | D |
| cmnm5Um | Gm |
| k2C | QtRNA |
| m1A | Um |
| m1G | Y |
| m1I | Ym |
| m2,2G | ac4C |
| m3C | cmnm5s2U |
| ms2i6A | cmo5U |
| o2yW | f5C |
| yW | galQtRNA |
| | gluQtRNA |
| | i6A |
| | m1Y |
| | m2A |
| | m2G |
| | m5C |
| | m5U |
| | m5Um |
| | m6A |
| | m6t6A |
| | m7G |
| | m8A |
| | mnm5U |
| | mnm5s2U |
| | s2C |
| | s4U |
| | t6A |
| | xA |
| | xG |
| | xU |

**Table S1- Detectable and undetectable RNA modifications.** Using tRNA sequencing data from E.coli, human, and mouse, we compared known modification positions (based on MODOMICS annotations) with signal of mismatches and RT failures. The modification types annotated ad "detectable modification" showed a clear mismatches/RT abortion signal in at least one position, at higher than 10% of the reads. "Undetectable modifications" did not show pattern of mismatches/RT failure at any annotated position

| U-C Codon box (avoidance of U ending) | Amino Acid | Main Gos |
|---|---|---|
| UUU/UUC | Phe | cell killing, antigen processing and presentation, canonical Wnt signaling pathway, T cell activation |
| UAU/UAC | Tyr | G protein-coupled receptor signaling pathway, brown fat cell differentiation, regulation of excretion, |
| UGU/UGC | Cys | regulation of renal system process, metal ion homeostasis, G protein-coupled receptor signaling pathway |
| GAU/GAC | Asp | positive regulation of small molecule metabolic process, L-amino acid transport, regeneration, canonical Wnt signaling pathway |
| CAU/CAC | His | oxidative DNA demethylation, negative regulation of cell fate commitment, heat generation |
| AGU/AGC | Ser (6-box) | regulation of carbohydrate catabolic process, acrosome reaction, DNA packaging, positive regulation of antigen processing and presentation |
| AAU/AAC | Asn | chromatin organization, fatty acid elongation, negative regulation of multicellular organismal process |

**Table S2- Go enrichment of genes avoiding the use of XXU vs XXC codons.**

We sorted genes based on the use of the indicated codons, and preformed GO enrichment analysis using Gorilla. The significant GO categories that were enriched in this analysis are presented here for each codon pair.

#### Supplementary Dataset Legends

**Dataset S1- GO categories expression in correlation with samples' 1st PC.** Correlation of expression of gene set belongs to the indicated GO categories with PC1, calculated based on active codon usage.

**Dataset S2- tRNA modification in activated T-cells. Summary of all positions with mismatches and RT-abortions in all samples.** Threshold for inclusion in the table is: minimum coverage of modified position = 20 reads, Modification fraction above 10%.

**Dataset S3- mRNA differential expression in activated T-cells.** Differential expression based on mRNA sequencing, as calculated by DEseq2. Marked in yellow are wybutosine biosynthesis enzymes.
